# Asymmetric ephaptic inhibition between compartmentalized olfactory receptor neurons

**DOI:** 10.1101/427252

**Authors:** Ye Zhang, Tin Ki Tsang, Eric A. Bushong, Li-An Chu, Ann-Shyn Chiang, Mark H. Ellisman, Jürgen Reingruber, Chih-Ying Su

**Affiliations:** Neurobiology Section, Division of Biological Sciences, University of California, San Diego, La Jolla, CA 92093, USA; National Center for Microscopy and Imaging Research, Center for Research in Biological Systems, University of California, San Diego, La Jolla, CA 92093, USA; Brain Research Center, National Tsing Hua University, Hsinchu 30013, Taiwan; Institut of Biology, École Normale Supérieure (IBENS), 46 rue d'Ulm, 75005 Paris, France; INSERM U1024, Paris, France

## Abstract

In the *Drosophila* antenna, different subtypes of olfactory receptor neurons (ORNs) housed in the same sensory hair (sensillum) can inhibit each other non-synaptically. However, the mechanisms underlying this unusual form of lateral inhibition remain unclear. Here we use recordings from pairs of sensilla impaled by the same tungsten electrode to prove that direct electrical (“ephaptic”) interactions mediate lateral inhibition between ORNs. Intriguingly, within individual sensilla, we find that ephaptic lateral inhibition is asymmetric such that one ORN exerts greater influence onto its neighbor. Serial block-face scanning electron microscopy of genetically identified ORNs and circuit modeling indicate that asymmetric lateral inhibition reflects a surprisingly simple mechanism: the physically larger ORN in a pair corresponds to the dominant neuron in ephaptic interactions. Thus, morphometric differences between compartmentalized ORNs account for highly specialized inhibitory interactions that govern information processing at the earliest stages of olfactory coding.

## INTRODUCTION

Ionic fluxes from neuronal activity lead to changes in the extracellular potential^1^, which can influence the excitability of adjacent neurons by electrical field effects, known as ephaptic interaction^2,3^. First observed between two axons brought together experimentally^4,5^, ephaptic interaction takes place between uninsulated neuronal processes packed into an electrically isolated microenvironment^2,3^. Such an arrangement commonly occurs in fascicles containing bundles of unmyelinated axons, such as the mammalian olfactory nerve^6^ and the interoceptive sensory system^7^, as well as in regions of the nervous system including the fish hindbrain, mammalian cerebellum, hippocampus and retina^1–3,8^. Despite their likely prevalence, field effects have long been considered nebulous^9^, as most neurons that are known to interact ephaptically also communicate via chemical synapses or gap junctions^2,3,8^. In addition, ephaptic interaction is notoriously difficult to study because it is enabled by high extracellular resistance and density of neural membranes^2,3^, none of which are amenable to *in vivo* experimental manipulation. Therefore, it remains unclear whether and how ephaptic interaction by itself is sufficient to influence circuit function.

Taking advantage of the powerful genetic toolkit of *D. melanogaster*, we showed that *Drosophila* olfactory receptor neurons (ORNs) housed in the same sensory hair, or sensillum, can inhibit each other^10^. Despite the lack of synaptic connections between neighboring ORNs, transient activation of one ORN rapidly suppresses the ongoing activity of its neighbor^10^. Electric circuit modeling suggested a potential mechanism for this non-synaptic signaling^10,11^: in the restrictive space of a sensillum lumen, the high resistance of the lymph^12^ favors the generation of field effects between compartmentalized ORNs^2,3^. However, whether ephaptic interaction underlies the inhibition between ORNs has not been directly demonstrated.

In addition, outstanding questions remain about the peripheral organization of ORNs. Most insect ORNs housed in the same sensillum exhibit distinct and characteristic extracellular spike amplitudes. Grouped ORNs are thus named “A”,“B” or “C” based on their relative spike amplitudes in descending order^13^. In fruitflies, olfactory sensilla contain up to four neurons, with the majority of them housing two ORNs^13,14^. Intriguingly, certain odorant receptors are only expressed in the large-spike “A” neurons, whereas others in the small-spike ORNs. For instance, in *D. melanogaster*, the Or22a receptor is expressed in the large-spike “A” neuron in the <u>a</u>ntennal <u>b</u>asiconic sensilla of type 3 (ab3A), which is paired with a small-spike neighbor expressing Or85b (ab3B)^15–17^. The ab3A(Or22a)-ab3B(Or85b) arrangement is also observed in other *Drosophila* species^18,19^. These evolutionarily conserved patterns of ORN arrangement point to functional constraints in neuronal organization. They also imply that grouped ORNs have distinguishable functional characteristics. However, beyond ligand specificity^20^, little is known about whether and how the large-spike “A” ORN functionally differs from its small-spike neighbor.

In this study, we use a novel experimental approach to provide direct experimental proof that ephaptic coupling alone is sufficient to drive lateral inhibition between ORNs. In addition, combining electrophysiological recordings, morphometric analysis based on serial block-face scanning electron microscopy (SBEM)^21^ and circuit modeling, we uncover a surprising functional disparity between compartmentalized ORNs in ephaptic inhibition and elucidate the underlying mechanism. Our study thus establishes the peripheral olfactory system of *Drosophila* as an ideal model to illuminate the impact of ephaptic interaction on circuit function and to determine its general operating principle.

## RESULTS

### Experimental proof of ephaptic inhibition between ORNs

How does one prove that ORNs inhibit each other ephaptically if the inhibition is not mediated by any protein or signaling molecule? In the absence of a manipulatable target, we addressed this question by testing whether direct electrical interaction is sufficient to cause inhibition between ORNs. If lateral inhibition proceeds electrically in a sensillum, by means of experimental manipulation, we expect to observe similar cross inhibition between ORNs housed in different yet electrically coupled sensilla. To test this possibility, we performed extracellular recordings using a metal electrode to connect the electric fields of two adjacent sensillum, a manipulation that allows for direct demonstration of the impact of electrical interation^22^.

In the control experiment examining an individual ab1 sensillum, the sustained spike response of ab1A to a prolonged dose of vinegar (large spikes in Fig. 1a) was markedly reduced by a pulse of superimposed CO_2_ that activated ab1C (small spikes). The inhibition of ab1A by CO_2_ was abolished in mutant flies lacking functional CO_2_ receptors^23^ (Fig. 1a), indicating that the inhibition of ab1A depends on the excitation of ab1C, consistent with our earlier results^10^.

**Figure 1.**
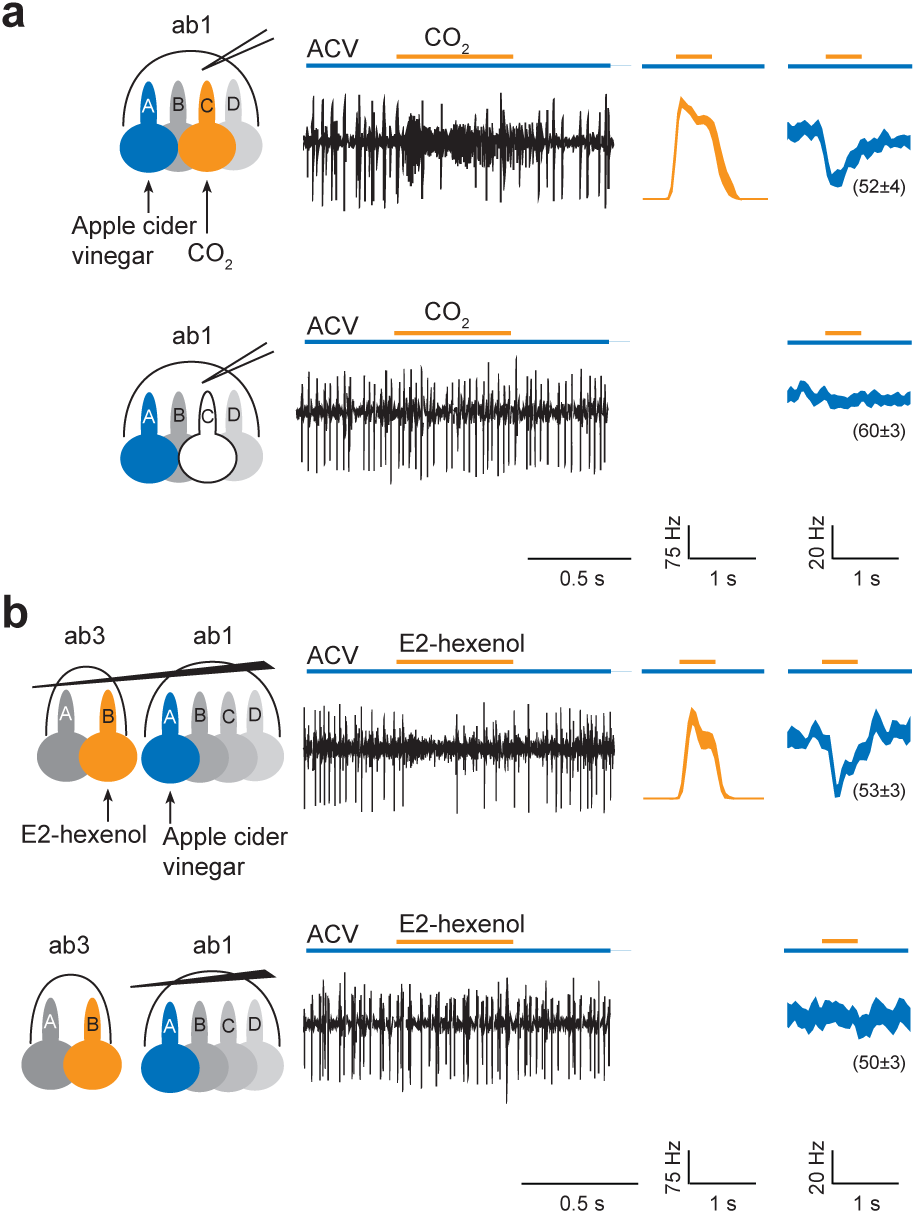
Direct electrical interaction drives lateral inhibition between ORNs. (**a**) The sustained response of ab1A was cross-inhibited by the transient activation of ab1C. Top: ab1A responded (large spikes in trace) to a sustained stimulus of apple cider vinegar (3x10^-3^ dilution, v/v in water, long blue bar). A 500-ms pulse of carbon dioxide (CO_2_, orange bar above trace) activated ab1C (small spikes). The response of ab1A was inhibited by the CO_2_ stimulus (decreased frequency of large spikes). In the average spike responses on the right, the orange trace represents the response of ab1C to CO_2_, and the blue trace represents the response of the large-spike neurons. ab1A and ab1B spikes could not be sorted reliably and were grouped. The sustained responses of the large-spike ORNs are indicated in the parentheses. Line width indicates s.e.m. Bottom: in the CO_2_ receptor mutant flies, CO_2_ did not activate ab1C or inhibit the sustained response of ab1A to vinegar. (**b**) Cross inhibition between electrically coupled ORNs. Top: a tungsten electrode was used to electrically couple two adjacent sensilla: ab3 and ab1. Activation of ab1A by a sustained stimulus of vinegar (3x10^-3^ dilution) was inhibited by the activation of ab3B housed in the electrically coupled sensillum by a pulse of E2-hexenol (10^-3^). Bottom: when the same two sensilla were no longer electrically coupled, E2-hexenol ceased to inhibit the sustained response of ab1A to vinegar. *n*=9.

Next we used a tungsten electrode to impale two adjacent sensilla, ab1 and ab3, in order to connect their electric fields so that the field changes in one sensillum can be detected by ORNs in the other sensillum. This approach eliminates the influence of other possible non-synaptic mechanisms, such as inhibition mediated by shared odorant binding proteins^24^. Using this paradigm, we recorded ORN activity from both ab1 and ab3 sensilla simultaneously (Fig. 1b, top panel). As predicted, the sustained response of ab1A was similarly inhibited by a superimposed pulse of E2-hexenol that excited the electrically coupled ab3B. The cross inhibition depended on the electric coupling because when we withdrew the tungsten electrode from the ab3 sensillum, we also abolished the inhibition of ab1A by E2-hexenol (Fig. 1b, bottom panel). These results thereby provide direct proof that the sustained ab1A response was cross-inhibited by the excitation of ab3B via the interconnected electric fields. In other words, direct electrical interaction, or ephaptic coupling, is sufficient to drive lateral inhibition between ORNs that share the same electric field.

### Field response and ephaptic inhibition

We next asked whether the large-spike “A” ORNs are functionally distinguishable from their small-spike neighbors in ephaptic interaction. In earlier work, we showed that lateral inhibition between ORNs is bidirectional: Transient activation of the “B” ORN inhibits the chronic response of the “A” neuron and vice versa^10^. However, it is unclear whether the bidirectional inhibition is equal in strength. To compare directly the strength of A-to-B and B-to-A inhibition is technically challenging because high frequency firing of the large-spike “A” neuron occludes spike activity of the “B” neuron. In addition, the degree of inhibition is influenced by the activity level of both ORNs^10^. To overcome these limitations, we sought to define the strength of ephaptic inhibition by other means.

According to an electric circuit model of a sensillum, neighboring ORNs shared the same electric field, also known as the transepithelial potential, which provides the driving force for odor-induced transduction currents^11^. As such, activation of one ORN reduces the transepithelial potential, thereby shunting currents away from its neighbor to cause ephaptic inhibition^10^. Thus, the degree to which activation of one ORN reduces the transepithelial potential, which can be measured as a change in the local field potential (LFP), indicates the magnitude of its ephaptic influence. The larger the LFP responses of one ORN, the more it can inhibit its neighbor.

To test this, we first identified odorants that strongly and selectively activate only one of the grouped ORNs (henceforth referred to as “private odorants”, Supplementary Fig. 1 and Supplementary Table 1). Using private odorants for the ab2 ORNs (ab2A: methyl acetate; ab2B: ethyl 3-hydroxy butyrate), we recorded LFP responses to 0.5-sec pulses of the private odorants, delivered either as individuals or as concurrent binary odor mixtures (Supplementary Fig. 2a). If there is no ephaptic inhibition, the LFP response to a binary odor mixture is expected to be the linear sum of the responses to its constituents. Thus, the difference between the linear sum and the measured LFP response indicates the magnitude of ephaptic inhibition.

As expected, we observed bidirectional inhibition using binary odor mixtures. Concurrent activation of ab2B by ethyl 3-hydroxy butyrate attenuated the LFP responses of ab2A to methyl acetate, and so did activation of ab2A to ab2B responses. Importantly, the degree of inhibition increased with higher LFP responses (Supplementary Fig. 2). Henceforth, we measured the LFP responses of an ORN to evaluate its ephaptic influence on its neighbor.

### Grouped ORNs of distinct spike amplitudes have different maximal field responses

To compare the LFP responses of the “A” and “B” ORNs, we first focused on the ab2 ORNs, for which multiple private odorants are available (Supplementary Table 1). When ab2A was stimulated by increasing concentrations of a private odorant, methyl acetate, its LFP responses plateaued at ~23 mV. In comparison, the near-saturated LFP responses elicited by ethyl 3-hydroxy butyrate, a private odorant for ab2B, were markedly smaller, only ~10 mV (Supplementary Fig. 2a1). Importantly, the LFP amplitudes were characteristics of the ORNs, regardless of the position of the electrode along the sensillum (not shown). We note that ethyl 3-hydoxy butyrate activates ab2B strongly and effectively; at 3x10^-4^ dilution, the odorant elicited a high spike response in ab2B (~250 spikes/sec), comparable to the spike responses of ab2A to methyl acetate (Supplementary Fig. 3 and Supplementary Table 1). Thus, the difference in the LFP responses in ab2A and ab2B is unlikely to have originated from different efficacies of the odorants. To verify this interpretation, we tested another pair of private odorants for the ab2 ORNs (ab2A: ethyl acetate; ab2B: E3-hexenol). We found that the near-saturated LFP response of ab2A remained markedly larger than that of ab2B (Fig. 2a1). These results indicate that strong activation of ab2A can reduce the shared electric field more than that of ab2B. In this context, ab2A is the dominant ORN in the pair.

**Figure 2.**
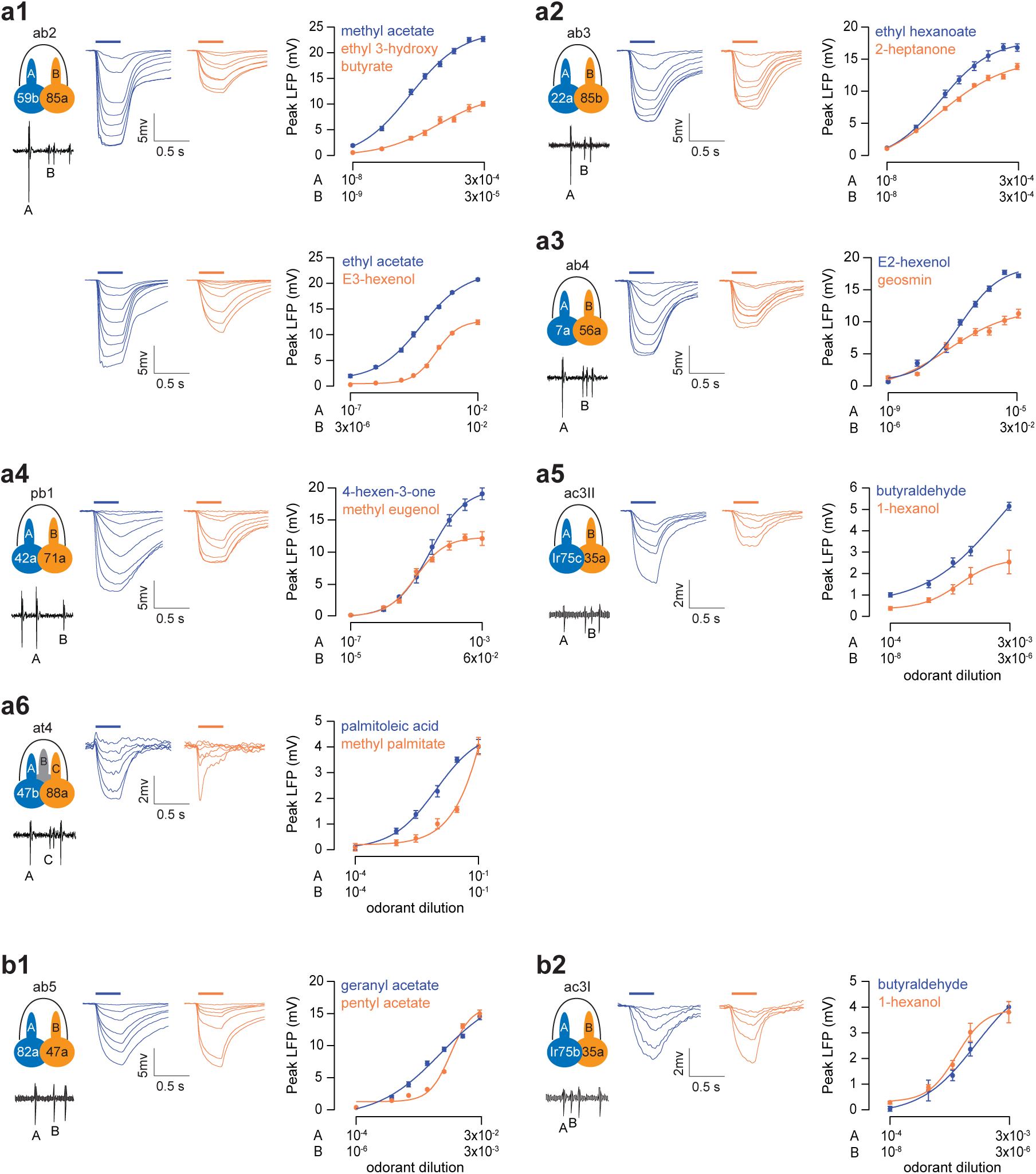
Comparison of the field responses of grouped neurons. (**a**) Dose-response relationships of grouped ORNs that have distinct extracellular spike amplitudes. (**a1**) Left: Spontaneous activity of ab2A (large spike) and ab2B (small spike). Middle: Average local field potential (LFP) responses of ab2A (blue traces) and ab2B (orange traces). Paired ORNs in the same sensilla were recorded in response to their respective private odorants at increasing concentrations. Right: Dose-response relationships of ab2A (blue) and ab2B neurons (orange) to methyl acetate and ethyl 3-hydroxy butyrate (top) or ethyl acetate and E3-hexenol, respectively (bottom). The absolute values of the peak LFP responses are shown. The highest and lowest concentrations of the “A” and “B” odorants are indicated logarithmically on the x-axis and aligned arbitrarily to facilitate comparison. (**a2-a6**) Additional sensillum types (ab3, ab4, pb1, ac3II and at4) were examined in a similar manner, as shown in (**a1**). (**b**) Average LFP responses and dose-response curves of grouped ORNs that exhibit similar extracellular spike amplitudes (**b1**: ab5; **b2**: ac3I). *n*=9 pairs of ORNs, except for ac3: *n*=6 pairs, mean ± s.e.m. Fit is with the Hill equation.

To test whether other large-spike “A” neurons also exert greater ephaptic influence upon their small-spike neighbors, we extended our analysis to additional sensillum types, selected on the basis of whether private odorants are available (Supplementary Table 1). In total, our analysis included 8 sensilla covering the major sensillum classes in the antenna and maxillary palp: large basiconic (ab2 & ab3), small basiconic (ab4 & ab5), coeloconic (ac3I & ac3II), trichoid (at4), and palp basiconic (pb1)^13,25,26^.

In 5 out of the 8 sensillum types examined, grouped ORNs exhibited notably different maximal LFP responses, including ORNs housed in the ab2, ab3, ab4, pb1, and ac3II sensilla. In each case where near-saturated responses were observed, the large-spike “A” neuron showed greater maximal LFP responses than its small-spike neighbor (Fig. 2a1–5). Analysis of the at4 ORNs could not be completed because their private odorants failed to elicit near-saturated LFP responses even at a high concentration (10^-1^) (Fig. 2a6, see Online Methods for details).

We note that a small minority of *Drosophila* olfactory sensilla contain ORNs of similar spike amplitudes, such as the ab5 and ac3I sensilla (Supplementary Fig. 4). In both cases, the near-saturated LFP responses of the grouped ORNs were nearly indistinguishable (Fig. 2b), suggesting that the neurons exert comparable ephaptic influence upon each other. Overall, our results indicate that when grouped ORNs exhibit markedly different spike amplitudes, they are more likely to exert unequal ephaptic influence upon each other, and most large-spike “A” neurons are the dominant neurons in this context.

### Grouped ORNs of distinct spike amplitudes have different tendencies to change spike responses

In addition to the strength of ephaptic influence, we compared the susceptibility of grouped ORNs to ephaptic influence. To this end, we analyzed the relationship between the peak spiking rate and the peak LFP amplitude of an olfactory response. The spike/LFP ratio of an ORN is a measure of the degree to which the neuron alters its spiking rate in relation to changes in its LFP response. For an ORN with a higher spike/LFP ratio, the neuron will experience a greater reduction in its spike response when the shared driving force is diminished by the activation of its neighbor. The ORN will therefore be more susceptible to ephaptic influence.

Upon stimulation, the frequency of ORN spiking is determined by the depolarization of the soma, which is the product of the transduction current and somatic input resistance (see Online Methods, modeling section). Although the transduction current cannot be measured directly in an *in vivo* preparation, it is thought to give rise to the LFP responses^27–29^. Hence, we used the LFP responses as a proxy for transduction currents in the following analysis.

We first stimulated ab3A and ab3B with 0.5-sec pulses of their respective private odorants at increasing concentrations and simultaneously recorded their LFP and spike responses (Fig. 3a1). By plotting the peak spike rate as a function of the peak LFP amplitude (absolute value), we found that the spike/LFP ratio was significantly higher for ab3B (Fig. 3a2), indicating that ab3B is more susceptible to ephaptic influence than ab3A. We note that the spiking rate of an insect ORN is determined not only by the amplitude but also by the kinetics of the LFP^29^. Thus, the variability in LFP kinetics introduced by variable transduction kinetics or odorant dynamics^29,30^ could have confounded our spike/LFP analysis.

**Figure 3.**
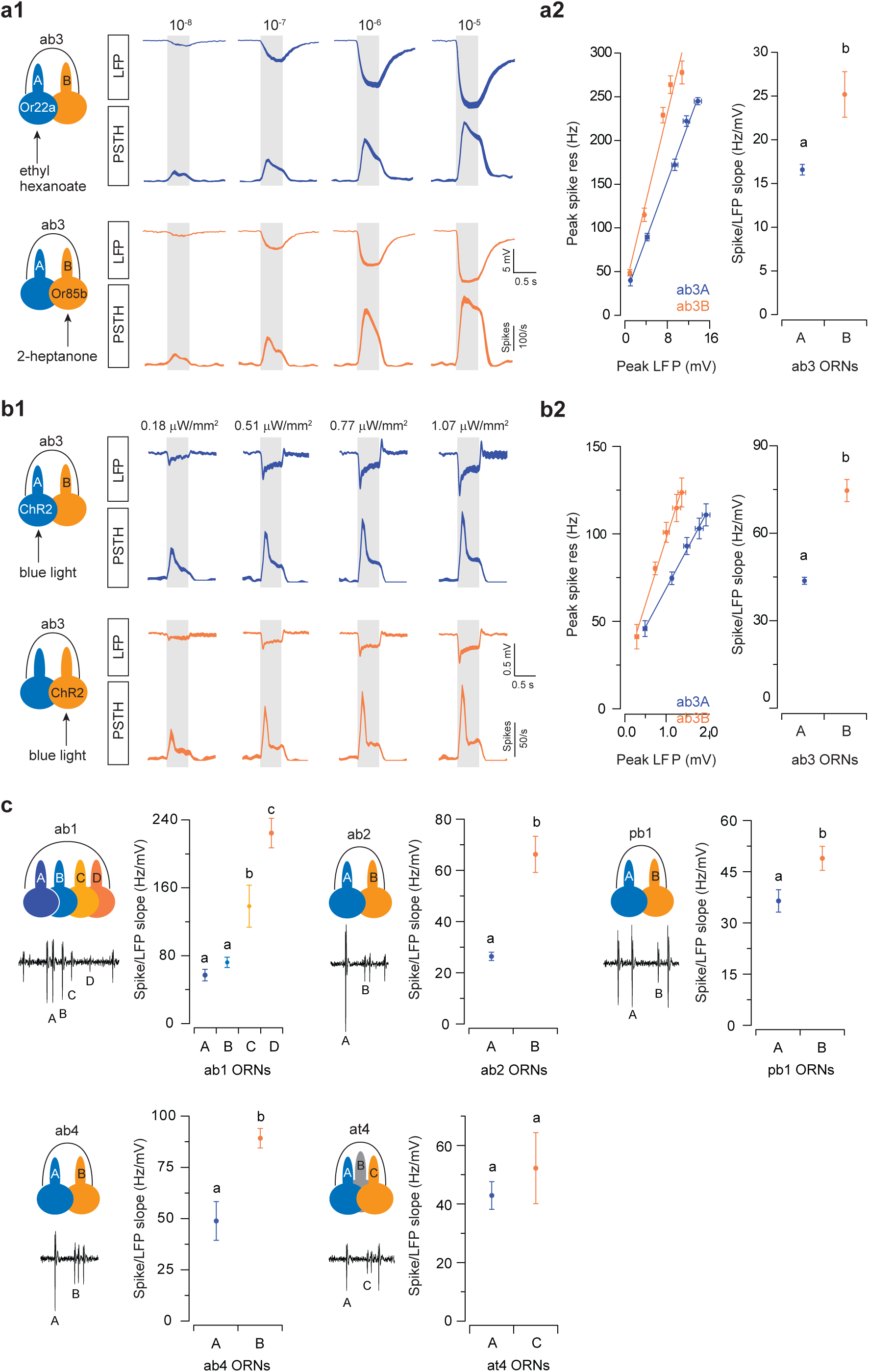
Comparison of the spiking properties of grouped neurons. (**a**) ab3 ORNs were selectively activated by their respective private odorants. (**a1**) Average LFP responses and the corresponding spike responses are shown for ab3A (blue) and ab3B (orange). (**a2**) Peak spike responses are plotted as a function of peak LFP responses (left). The absolute values of the peak LFP responses were used for all analyses. Lines indicate linear fits (*y = ax + b*). *n*=9 pairs of ORNs, mean ± s.e.m. The respective “*a”* coefficients (spike/LFP slope) are plotted for comparison (right). Error bars = s.d. (**b**) Similar to (**a**) except that ab3A and ab3B were activated optogenetically. H134R-Channelrhodopsin2 (ChR2) was expressed in either ab3A or ab3B by the GAL4-UAS system. ORNs were activated by 500-ms pulses of blue light (470 nm, irradiances are indicated above). *n*=9, parallel experiments. (**c**) Optogenetic analyses in five additional sensilla. ORNs expressing H134R-ChR2 were activated by blue light and the peak spike and LFP responses were analyzed as shown in (**b**). Error bars = s.d. *n*=9, parallel experiments. Statistical analysis was performed with ANCOVA and significant differences (*P* < 0.05) are denoted by different letters.

To address this concern, we used an optogenetics approach so that identical stimuli can be used to activate grouped ORNs. When light activated either ab3A or ab3B expressing Channelrhodopsin2 (H134R-ChR2)^31^, the LFP responses increased with stimulus intensity (Fig. 3b1). Consistent with our earlier analysis with odor stimulation, the spike/LFP ratio was also significantly higher for ab3B in the optogenetic assay (Fig. 3b2). Of note, a similar spike/LFP relationship between ab3A and ab3B was observed even with a lower level of functional H134R-ChR2, achieved by lowering the concentration of the chromophore (all *trans*-retinal) fed to the flies (Supplementary Fig. 5). Therefore, the difference in the spike/LFP ratios between ab3A and ab3B was unlikely to have been affected by the exact expression level of H134R-ChR2.

We then extended our optogenetic analysis to 5 additional sensillum types, including ab1, ab2, ab4, pb1 and at4. In each case, except for the at4 ORNs of which the spike amplitudes were not as distinct (Supplementary Fig. 4), the small-spike ORNs exhibited significantly higher spike/LFP ratios than the neighboring “A” ORNs (Fig. 3c). Notably, even in the ab1 sensillum that houses four ORNs, the relative spike amplitudes of the neurons (ab1A ≥ ab1B > ab1C > ab1D) remained indicative of the rank order of the spike/LFP ratios (ab1A ≥ ab1B > ab1C > ab1D) (Fig. 3c).

To test whether grouped ORNs with similar spike amplitudes have similar spike/LFP ratios, we examined ab5 and ac3I, the two sensilla which house ORNs of similar spike amplitudes (Supplementary Fig. 4). For technical reasons, we could not perform the optogenetic analysis in these sensilla (see Online Methods for details). Therefore, we ectopically expressed Or83c, an odorant receptor tuned to farnesol^32^, in either ab5A or ab5B in order to compare their responses to identical stimuli. Farnesol does not activate the cognate receptors of the ab5 ORNs^32^. Analyses of farnesol-induced responses indicated that ab5A and ab5B indeed have similar spike/LFP relationships (Supplementary Fig. 6a). Consistent with this result, analyses with private odorants for ab5 and ac3I ORNs also suggested that the neighboring neurons exhibit similar spike/LFP relationships (Supplementary Fig. 6b,c).

Taken together, our odorant and optogenetic analyses show that in most sensillum types, ephaptic interactions are asymmetric in that the large-spike “A” neuron is dominant. This dominance is due to (1) the “A” neuron’s greater ability to reduce the shared driving force (measured as LFP responses, Fig. 2), and (2) its lower susceptibility to ephaptic influence, evaluated by its lower propensity to change spiking rate in response to changes in LFP (Fig. 3).

### Overexpression of odorant receptors does not enhance the LFP response of a “B” ORN

What determines the difference in the maximal LFP responses of the “A” and “B” ORNs? One possibility is that the density of odorant receptors expressed in the “B” ORNs is typically lower than in the “A” ORNs. As such, fewer receptors in the “B” neuron can be activated by odor stimulation, resulting in a smaller increase in the transduction conductance and hence a smaller LFP response. To test this possibility, we used the GAL4/UAS system to overexpress the odorant receptor, Or85a, in its cognate ab2B ORNs. To ensure dendritic targeting of Or85a, we also overexpressed Orco, an obligatory co-receptor required for OR targeting and function^33^. Immunostaining of the epitope-tagged odorant receptors indicated successful dendritic targeting (Supplementary Fig. 7).

In the wildtype control, strong activation of ab2A resulted in a large LFP response (~28 mV), markedly higher than the near-saturated LFP response of ab2B (~12 mV) (Fig. 4a1). Interestingly, in flies overexpressing Or85a/Orco in ab2B, the maximal ab2B LFP response remained similar to that of the control, well below the ab2A counterpart (Fig. 4a2). That is, overexpression of Or85a did not increase the LFP responses of ab2B. A plausible scenario is that the expression level of endogenous Or85a is already high, likely close to saturation, such that overexpressed Or85a only replaces the endogenous receptor in the sensory dendrite without further increasing its receptor density to elevate the LFP responses. Thus, the smaller, near-saturated LFP response of a “B” ORN is unlikely to have arisen from a lower receptor density.

**Figure 4.**
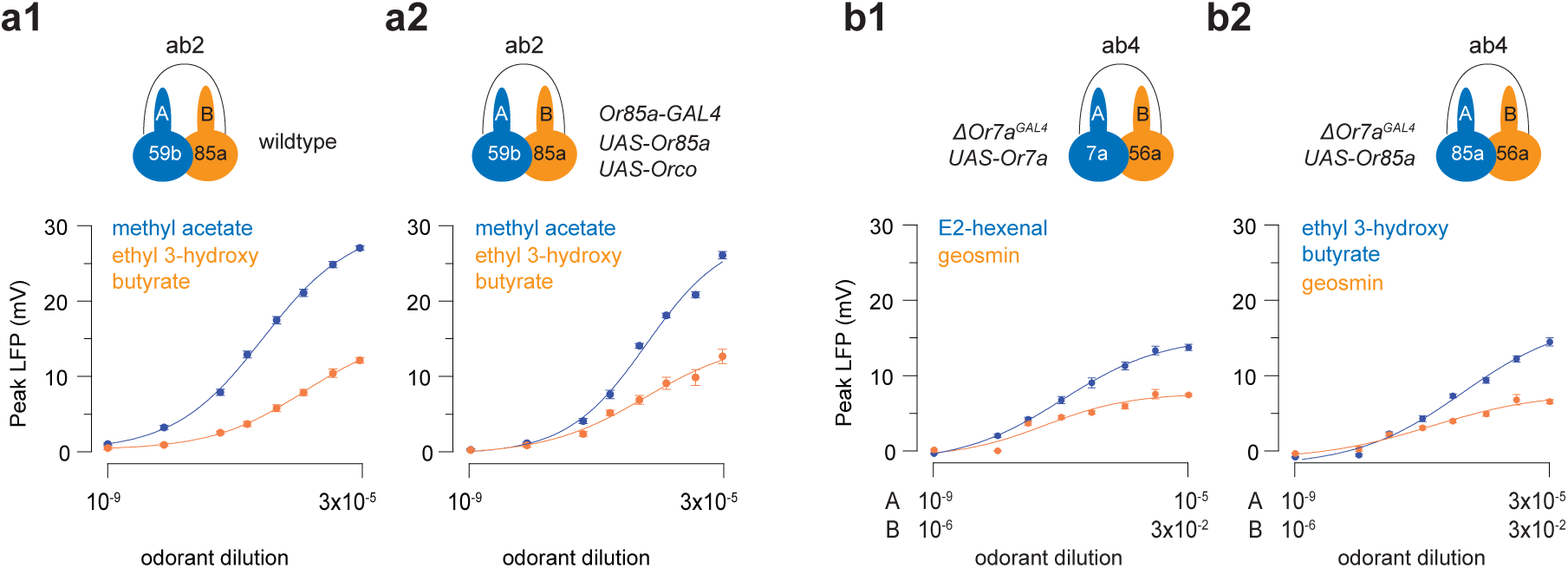
Overexpressing or swapping odorant receptors does not change the maximal LFP responses of an ORN. (**a**) Receptor overexpression in the ab2B ORNs. ab2A responded to methyl acetate (top traces, large spikes) and ab2B to ethyl 3-hydroxybutyrate (bottom traces, small spikes). (**a1**) In control flies, ab2A had a greater near-saturated LFP response than ab2B (similar results observed in Fig. 2a1). (**a2**) Overexpression of the odorant receptor complex, Or85a/Orco, in ab2B did not increase its maximal LFP responses. (**b**) Receptor swap in the ab4A ORNs. (**b1**) Genetic rescue of Or7a expression in the *ΔOr7a*^Gal4^ mutant flies (Δab4A:Or7a) restored the responses to E2-hexenal (top trace, large spikes). The neighboring neuron, ab4B, responded to geosmin (bottom traces, small spikes). The rescued ab4A ORNs had a greater near-saturated LFP response than ab4B, as in wildtype flies (see Fig. 2a3). The highest and lowest concentrations of the “A” and “B” odorants are indicated logarithmically on the x-axis and aligned arbitrarily to facilitate comparison. (**b2**) Ectopic expression of Or85a, the cognate receptor of ab2B, in Δab4A resulted in a similar near-saturated LFP response as in Δab4A:Or7a. *n*=9 pairs of ORNs, mean ± s.e.m. All results in (**a**) or (**b**) are from parallel experiments.

### Swapping odorant receptors does not lower the LFP response of an “A” ORN

Next, we examined the contribution of receptor identity to the maximal LFP responses. In a sensillum, neighboring ORNs express different odorant receptors^16,34,35^. Given that *Drosophila* odorant receptors are ligand-gated cation channels^36,37^, it is possible that the conductance of the receptors expressed in the “A” ORNs is typically larger than in the “B” neurons. If so, when the cognate receptor of an “A” ORN is replaced by a “B” neuron receptor, one would expect a marked reduction in the maximal LFP response of the “A” neuron.

To test this possibility, we performed receptor-swap experiments in the ab4A ORNs, of which the cognate receptor is Or7a^15^. We chose ab4A because Or7a receptor mutants (*ΔOr7a*^GAL4^) are readily available^38^, and because its small-spike neighbor, ab4B, is narrowly tuned to geosmin^39^. In *ΔOr7a*^GAL4^ mutants, the response of ab4A (Δab4A) to an Or7a ligand, E2-hexenal, is completely abolished (not shown). Genetic rescue of Or7a in Δab4A (Δab4A:Or7a) restored the response; the restored LFP dosage curve was similar to that of the wildtype ab4A, above the ab4B curve (Fig. 4b1 and 2a3). When a “B” neuron receptor (Or85a/ab2B) was expressed in ab4A instead (Δab4A:Or85a), the LFP dosage curve to an Or85a ligand, ethyl 3-hydroxybutyrate (Fig. 4b2), was remarkably similar to that of Δab4A:Or7a to E2-hexenal (Fig. 4b1). These results suggest that the characteristic near-saturated LFP responses of an ORN are not influenced by receptor density or identity but likely by other ORN features.

### Morphometric analysis of grouped ORNs

What then underlies the asymmetry in ephaptic interactions? The size difference between grouped ORNs is likely a key. In most fly sensilla, grouped ORNs have differing sizes^40^, and the large-spike “A” neurons were postulated to have larger dendritic calibers^41–43^. However, without a genetically encoded EM marker, it was impossible to assign ORN identity. Notably, the ORN with a larger soma would have a smaller input resistance, which could account for the smaller spike/LFP ratio of the “A” neurons. In addition, a larger surface area of the sensory dendrites could give rise to a larger maximal LFP response, also characteristic of the “A” neurons. Therefore, we hypothesized that the dominant “A” neuron is larger than its neighbor.

To test this hypothesis, we endeavored to measure the morphometric features of genetically identified ORNs using electron microscopy (EM). To this end, we developed a method termed CryoChem, which allows for quality morphological preservation of genetically labeled structures for volume EM imaging^44^. With OR-specific drivers, we expressed an EM marker, APEX2 (enhanced <u>a</u>scorbate <u>pe</u>ro<u>x</u>idase 2)^45^, in select ORNs to render them electron dense through diaminobenzidine (DAB) labeling. As a proof of principle, we first generated 3D volumes of at4A (Or47b>APEX2) and at4C (Or88a>APEX2) using SBEM (Fig. 5a,b). Among the three at4 ORNs, we identified at4A and at4C as the largest and intermediate-sized neurons in the group, respectively, and used this information to assign neuronal identity to the unlabeled ORNs. By comparing the APEX2-labeled and unlabeled ORNs, we verified that APEX2/DAB labeling does not significantly alter the morphometric features of ORNs (Fig. 5c).

**Figure 5.**
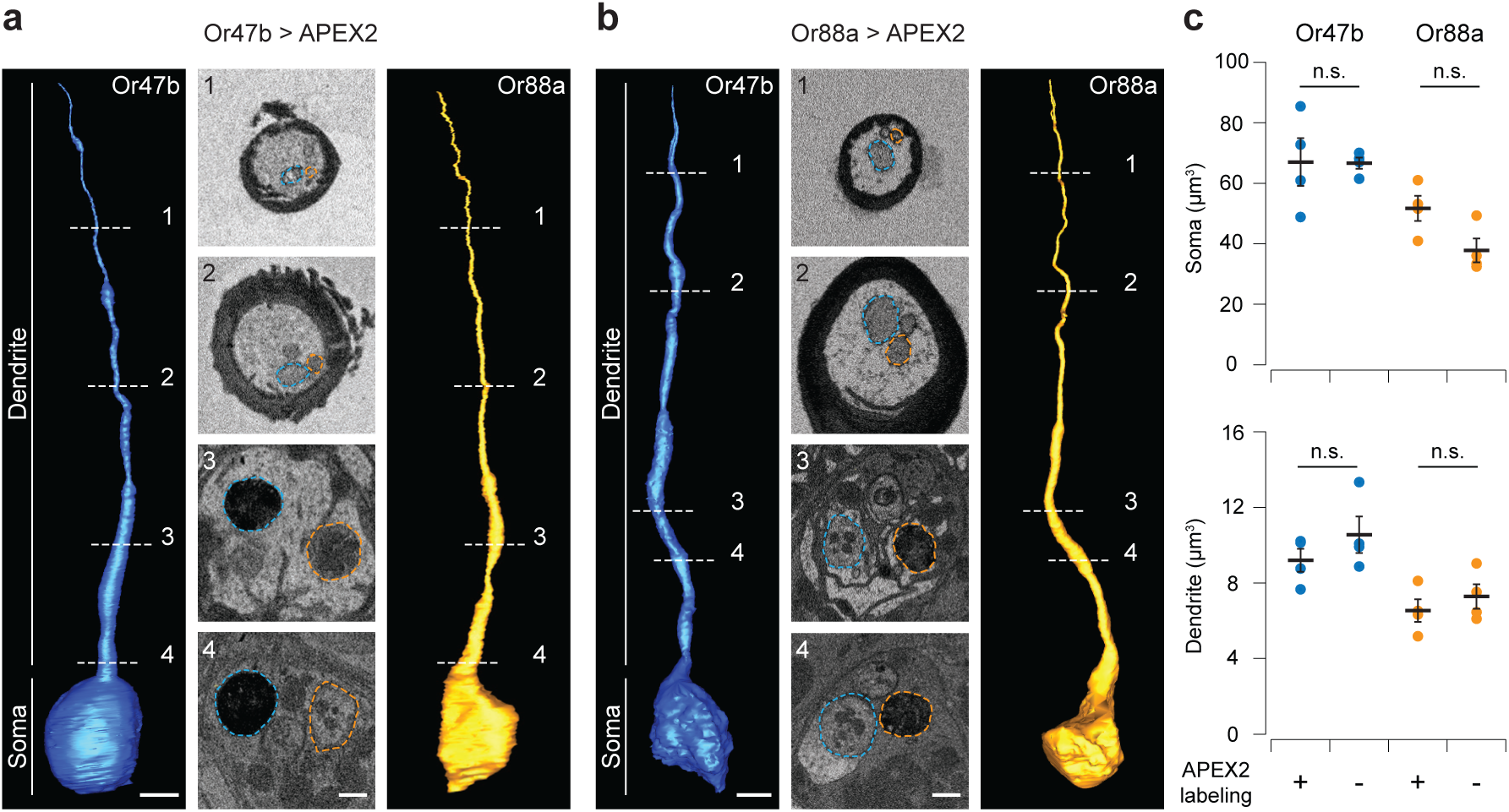
Morphometric measurements of genetically identified ORNs with SBEM. (**a-b**) 3D reconstruction of the Or47b (blue) and Or88a ORNs (orange) based on the SBEM volumes generated from Or47b>APEX2 and Or88a>APEX2 antenna, respectively. (**a**) Or47b ORNs expressing APEX2 were labeled with DAB. Sample SBEM images are shown in the middle panel. Dotted lines outline the Or47b ORN (blue) and its intermediate-sized neighbor (orange). (**b**) Or88a ORNs expressing APEX2 were labeled with DAB. In the sample SBEM images, dotted lines outline the Or88a ORN (orange) and its largest-sized neighbor (blue). Scale bars: 2 µm for 3D models, 500 nm for SBEM images. (**c**) The soma and dendritic volumes of the Or47b and Or88a ORNs, mean ± s.e.m. *n*=4 pairs of ORNs. APEX2-labeled vs. unlabeled Or47b soma, *P* = 0.725; Or88a soma, *P* = 0.224; Or47b dendrite, *P* = 0.274; Or88a dendrite, *P* = 0.167, *t*-test.

### Dominant “A” ORNs are physically larger than their neighbors

Next, we expanded our morphometric analysis to a total of 5 sensillum types (ab3, ab4, ac3II, at4 and ab5) that cover major sensillum classes. In the sensilla where grouped ORNs exhibit distinct spike amplitudes (ab3, ab4, ac3II and at4), the “A” ORNs were significantly larger than their small-spike neighbors with respect to the soma, inner and outer (sensory) dendrites (Fig. 6a-d, Supplementary Table 2). Notably, ephaptic interactions were also asymmetric in these sensilla (with the exception of at4 where the dose-response analysis was incomplete), supporting our hypothesis that morphometric disparity between grouped ORNs underlies the asymmetry in ephaptic interactions. To further test this hypothesis, we examined the ab5 ORNs, which exert equal ephaptic influence onto each other (Fig. 2 and Supplementary Fig. 6) and would thus be predicted to be of similar size. Indeed, in ab5A and ab5B we found that their somata, and likely also the sensory dendrites, were similar in size (Fig. 6e, Supplementary Table 2, see below for disparity in inner dendritic volumes).

**Figure 6.**
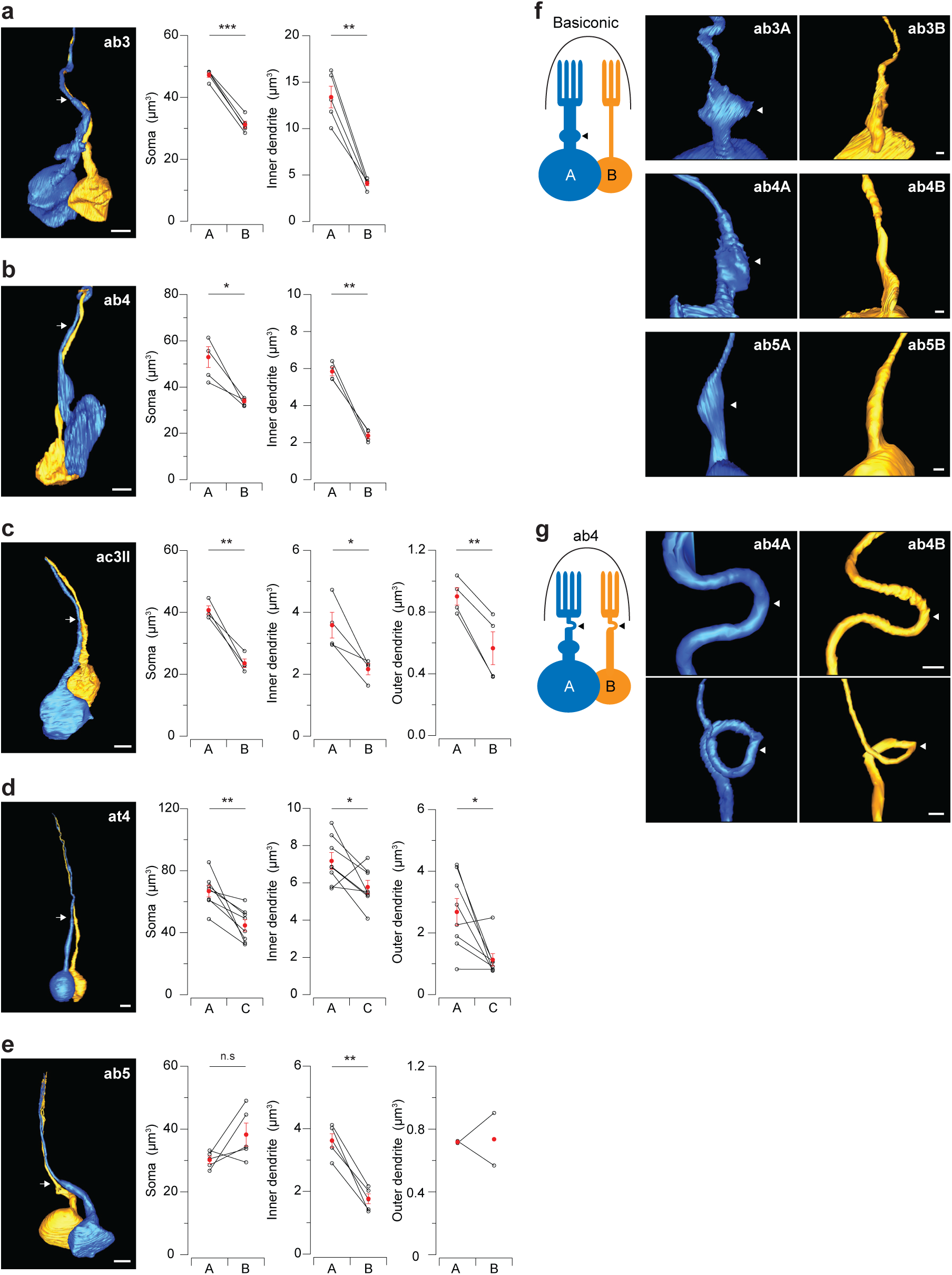
Systematic morphometric analysis of grouped ORNs. (**a-e**) Volumes of the soma, inner and outer dendrites of the paired ORNs in five sensillum types. (Left) Sample 3D reconstruction based on SBEM images. Arrows indicate the cilium base, a constricted region separating the inner and outer dendrites. Due to technical limitations, the outer dendrites of most basiconic sensilla could not be completely reconstructed (see Online Methods for details). Scale bars: 2 µm. Lines connect measurements from paired ORNs, mean ± s.e.m. *n*=4-5 for all except for at4, *n*=8. \**P* < 0.05, \*\**P* < 0.01, \*\*\**P* < 0.001, paired *t*-test. Error bars and statistics are unavailable for the ab5 outer dendrites as only two data points were obtained. (**f-g**) Novel morphological features. (**f**) An enlargement of the inner dendrite (arrow head) was observed in the “A” ORNs in all three characterized basiconic sensilla. (**g**) A bend or loop (arrow head) was observed in the unbranched outer dendrite of the ab4 ORNs. Scale bar: 500 nm.

Beyond morphometric measurements, our SBEM experiments also revealed novel ORN morphological features. For example, in the basiconic sensilla, but not in other sensillum types, we noticed a marked enlargement of the inner dendrite in every “A” ORN (Fig. 6f). This feature explains why the inner dendrite of ab5A is significantly larger than that of ab5B, despite their otherwise similar sizes (Fig. 6e). In addition, in the ab4 sensilla, we observed bending or looping of the unbranched outer dendrite around the base of the sensillum (Fig. 6g). These observations further highlight the diversity of ORN morphology within the same or among different sensillum types.

### An electric circuit model predicts the asymmetric ephaptic relationship between grouped ORNs

We used mathematical modeling to explain how the morphometric disparity between ORNs contributes to their asymmetric ephaptic interactions. The existing electric circuit model of a sensillum assumes that neighboring ORNs have identical passive electrotonic properties^11^. However, our morphometric analysis suggests otherwise. We therefore took into account that the input resistance of an ORN is inversely proportional to the surface area of the soma, and that the conductance change upon odor stimulation scales with the surface area of the sensory dendrite (Fig. 7a, see Online Methods for modeling details).

**Figure 7.**
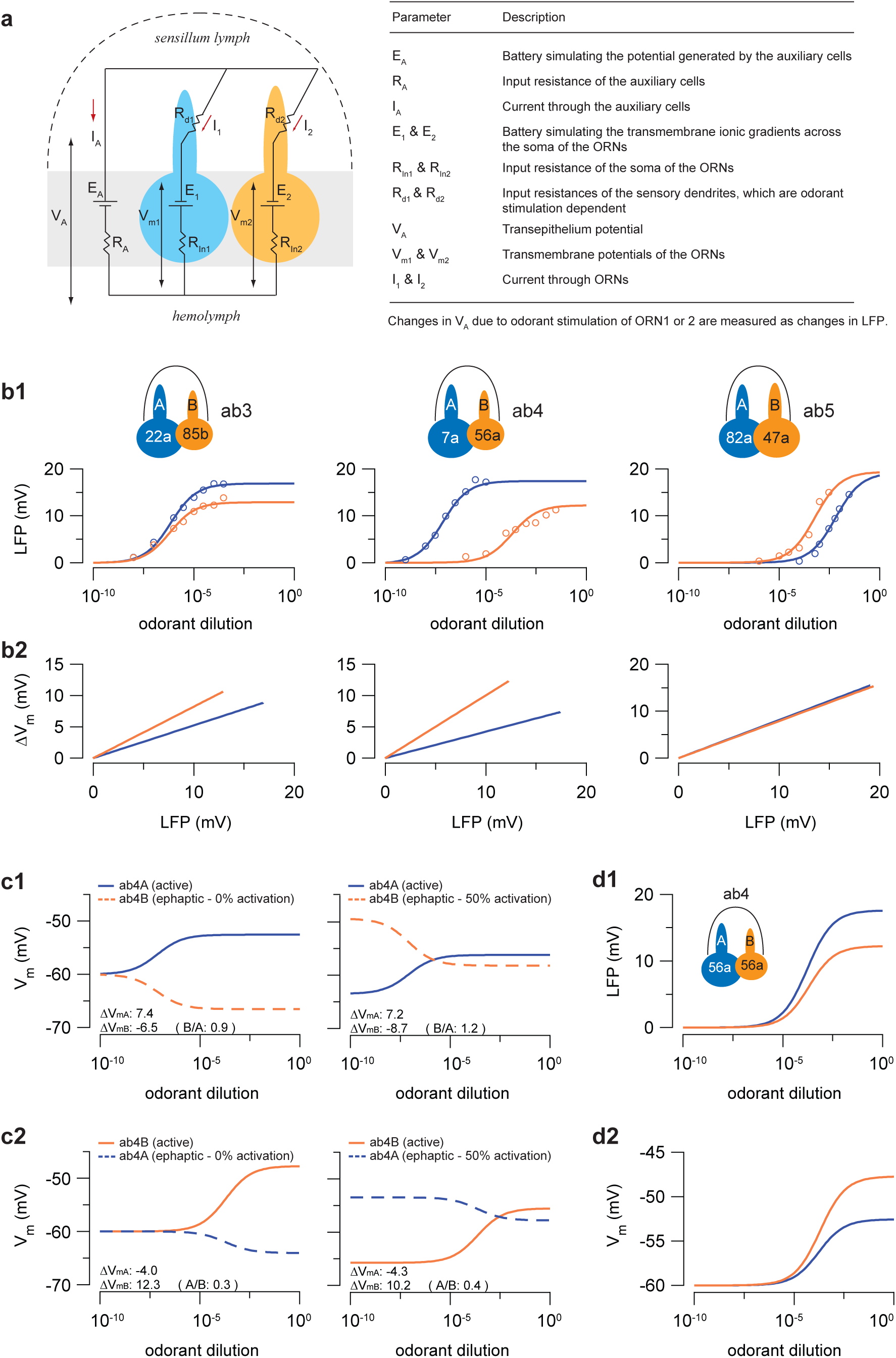
An electric circuit model for compartmentalized ORNs. (**a**) Passive electric circuit model of a sensillum consisting of two ORNs and an auxiliary cell (gray rectangle). (**b**) Fitting and simulation of ORN responses. (**b1**) Simultaneous fitting of the LFP responses of the ORNs housed in the ab3, ab4 and ab5 sensilla. Identical parameters were used for all ORNs, except for the morphometric parameters and odorant sensitivity, which are ORN-specific and determined based on the experimental data whenever available. Each curve describes the dose-response relationship of an ORN when activated by a private odorant. Empty circles indicate measured LFP responses. Odorant concentrations are plotted logarithmically on the x-axis. (**b2**) Simulation of ORN depolarization (ΔV_m_) in relation to the LFP response. (**c**) Simulation of the transmembrane potential of the ab4 ORNs (V_m_) when either ab4A (**c1**) or ab4B (**c2**) was activated by increasing concentrations of private odorants. Two different activation states of the neighboring ORN (0% and 50%) were considered. Depolarization of one ORN (“active”, solid line) ephaptically hyperpolarizes its neighbor (“ephaptic”, dashed line). ∆V_mA_ and ∆V_mB_: maximal change in V_m_ (mV) for ab4A and ab4B, respectively. A/B or B/A: the absolute value of ∆V_mA_/∆V_mB_ (A/B) or ∆V_mB_/∆V_mA_ (B/A). (**d**) Simulation of the LFP responses and transmembrane potential changes of the ab4 ORNs, assuming that both ORNs express the same odorant receptor, Or56a. Except for ab4A odorant sensitivity, all fitting parameters are as in (**b1**) for ab4. Blue: large-spike “A” ORN. Orange: small-spike “B” ORN. See Online Methods for details on modeling, data fitting and parameters.

In our circuit model, the ORN-specific parameters are the odorant sensitivity and the surface areas of soma and sensory dendrite. For the remaining modeling parameters, such as the resting membrane potential and membrane resistivity, identical values were assumed for all ORNs. First we evaluated our model by fitting the peak LFP responses of the ORNs housed in the ab3, ab4 and ab5 sensilla, of which electrophysiology and morphometric data were both available. We fitted the responses from the three ORN pairs at the same time with identical free parameters using the *Data2dynamics* framework with a nonlinear least-squares optimization algorithm^46^. The fitted dose-response curves indicated that our circuit model well describes the experimental data (Fig. 7b1). They also confirmed that the larger sizes of the “A” neurons result in higher maximal LFP responses, as evidenced in the fittings of the ab3 and ab4 ORNs, which are of different sizes, but not the ab5 ORNs, which are of similar sizes (Fig. 7b1). For the ab3 and ab4 sensilla, where measurements for the sensory dendrites were unavailable, the fitted parameters indicate that the dendritic surface areas of ab3A and ab4A are both larger than those of their respective neighbors (Online Methods). We also used the model to predict the relationship between ORN depolarization (ΔV_m_) and LFP responses (Fig. 7b2). Given that the peak spike response of an ORN is a function of ∆V*_m_*^29^, the smaller ∆V_m_/LFP slopes of the “A” neurons are consistent with their smaller spike/LFP ratios (Fig. 3).

Our experimental data predict that ephaptic interaction is asymmetric between grouped ORNs in most sensilla (Fig. 2–3). To test whether our model describes asymmetric inhibition, we simulated the transmembrane potential of both ab4 ORNs when either ab4A or ab4B was activated by a private odorant at increasing concentrations. According to our model, depolarization of one ORN hyperpolarizes its neighbor (Eqs. 11 & 12, Online Methods). We observed this relationship when both ORNs were assumed to be initially inactive (Fig. 7c, 0% activation) and when one ORN was assumed to be chronically activated by a background odorant, a paradigm used in Fig. 1 and in our earlier work^10^ (Fig. 7c, 50% activation). We then evaluated the strength of ephaptic inhibition between neighboring ORNs by comparing the degree to which depolarization of one ORN hyperpolarizes its neighbor. A higher value of the ratio (hyperpolarization/depolarization) indicates stronger ephaptic inhibition. By comparing the absolute values of ∆V_mB_/∆V_mA_ (Fig. 7c1) and ∆V_mA_/∆V_mB_ (Fig. 7c2), we found that activation of ab4A was indeed more effective in inhibiting ab4B than the reverse situation, indicating that A-to-B ephaptic inhibition is stronger than B-to-A inhibition.

In our model, the only parameters that are different between grouped ORNs are the neuronal size and odorant sensitivity, both of which can in principle contribute to asymmetric ephaptic interaction. To illustrate the impact of neuronal size, we simulated ab4 ORN responses in a hypothetical situation where both ORNs express the same odorant receptor, Or56a. Even with identical odorant sensitivity, the larger size of ab4A was sufficient to yield a larger maximal LFP response (Fig. 7d1) and a smaller increase in the transmembrane potential (Fig. 7d2). In other words, morphometric differences alone can account for the asymmetric ephaptic interactions between grouped ORNs.

## DISCUSSION

We demonstrate here that in the absence of any synaptic communication, ephaptic interaction is sufficient to influence circuit function by driving lateral inhibition between compartmentalized ORNs to markedly reduce neuronal spike activity. Surprisingly, we find that across sensillum types, most large-spike “A” neurons can exert greater ephaptic influence onto their neighbors and are also less susceptible to ephaptic influence. Mechanistically, the functional disparity arises from morphometric differences between compartmentalized ORNs. Together, our study describes a highly specialized inhibitory interaction that governs information processing at the earliest stages of olfactory coding. It also establishes the peripheral olfactory system of *Drosophila* as a model to understand the impact of electric field effects on neural circuit function.

Ephaptic interactions are asymmetric between grouped ORNs with distinct spike amplitudes. Conversely, grouped ORNs exhibiting similar spike amplitudes exert similar ephaptic influence on each other (Fig. 2b and Supplementary Fig. 6). This relationship suggests that the extracellular spike amplitude of an ORN and its ephaptic influence are regulated by a common factor. Indeed, as with ephaptic influence, the extracellular spike amplitude of a neuron also negatively correlates with its input resistance^47^. Thus, the larger ORN in a pair, which has the smaller input resistance, is expected to have larger spike amplitude as well as ephaptic dominance over its neighbor.

In addition to fly ORNs, there are reports of asymmetric inhibition between other primary sensory neurons. For example, in fly vision, R7 and R8 photoreceptors of the same ommatidium inhibit each other synaptically. Interestingly, the mutual inhibition between R7 and R8 is also asymmetric^48^. The asymmetry may have arisen from the unequal numbers of reciprocal synapses^49^. In the bumblebee galea sensilla, gap junction-mediated inhibitory coupling is unequal between compartmentalized gustatory receptor neurons^50^. Therefore, although the precise mechanisms may vary, asymmetric lateral inhibition between adjacent primary sensory neurons may represent a conserved computational motif whereby sensory inputs are unequally processed at the periphery before being transmitted to higher brain centers.

Ephaptic interactions between ORNs allow odor-mixture information to be processed by the first neurons of an olfactory circuit^10^. To evaluate the extent to which natural odors activate grouped ORNs simultaneously, we surveyed volatile compounds from several food sources for fruit flies (Supplementary Table 3). From 16 fruits and fermented foods, we identified 51 odorants that can excite at least one ORN with responses ≥ 50 spike/s, based on published datasets^14,20,39^. Among them, 30 individual odorants were capable of co-activating neighboring ORNs. For instance, ethyl hexanoate elicited significant responses in both ab3A and ab3B ORNs (Supplementary Table 3). Strikingly, all of the analyzed odor sources contained volatiles that excited at least one pair of the grouped ORNs, suggesting that grouped fly ORNs are commonly co-activated by natural odors.

What then is the significance of asymmetric lateral inhibition between grouped ORNs? In such a situation, this operation may provide a peripheral mechanism for evaluating countervailing signals and favoring the propagation of the input carried by the large-spike ORNs. In a mixture, odorants that excite the small-spike ORNs are more likely to be masked by odorants activating their large-spike neighbors. In support of this idea, we previously found that activation of ab1A by vinegar odors attenuated the aversiveness of CO_2_ detected by ab1C^10^. In future studies, it will be important to determine how odorants that excite the large-spike ORNs qualitatively differ from those that excite the small-spike neurons.

## Acknowledgements

We thank Jeffry Isaacson, Johannes Reisert, Jing Wang, Larry Squire, Gareth Thomas and Renny Ng for comments on the manuscript; Szu-Ying Chen, Vivian Nguyen, Renny Ng, Edie Zhang, Martin Orden, Uma Talagadadivi, Ben Damasco, Tiffany Tsai and Cesar Nava-gonzales for assistance in ORN segmentations; Cesar Nava-gonzales for assistance in the analysis of natural odors. This work was supported by Frontiers of Innovation Scholars Program and Croucher Foundation Scholarship to T.K.T.; Hellman Fellowship, the Ray Thomas Edwards Foundation Career Development Award, Kavli Institute for Brain and Mind Innovative Research Grant and NIH R01DC015519 to C-Y.S. National Center for Microscopy and Imaging Research (NCMIR) technologies and instrumentation are supported by grant GM103412 from the National Institute of General Medical Sciences.

## Author Contributions

Y.Z., T.K.T. and C-Y.S. designed the experiments. Y.Z. performed sensillum recordings, data analysis and natural odor analysis. T.K.T. and E.B. performed SBEM imaging with support from M.E. T.K.T. performed immunostaining and morphometric analysis. L-A.C conducted preliminary morphometric analysis using super-resolution confocal microscopy with support from A-S.C. J.R. performed computational modelling. Y.Z., T.K.T., J.R. and C-Y.S. wrote the manuscript with inputs from all authors.

## ONLINE METHODS

***Drosophila* stocks**. Flies were raised on standard cornmeal medium at 25°C, ~60% relative humidity in an incubator with a 12-hr light/dark cycle. Female CS flies 5–7 days post eclosion were used in all experiments unless noted otherwise. For the ablation (*UAS-rpr*) and optogenetic (*UAS-H134R-ChR2*) experiments, 5-day old females were used; for the SBEM (*UAS-APEX2*) experiments, 6–8 day old females were used. For further information on genotypes, refer to Supplementary Table 4.

**Single-sensillum recordings**. A fly was wedged into the narrow end of a truncated plastic 200-μl pipette tip to expose the antenna, which was subsequently stabilized between a tapered glass microcapillary tube and a coverslip covered with double-sided type^51^. Single-unit recordings were performed essentially as described10. Briefly, electrical activity of the ORNs was recorded extracellularly by placing a sharp electrode filled with 0.6X sensillum lymph Ringer solution28 into a sensillum and the reference electrode filled with the same Ringer solution was placed in the eye or the clypeus (for at4 recordings). For recordings performed with a tungsten electrode, a tungsten rod (0.01 × 3 inch, 717000, A-M Systems) secured in an electrode holder (ST50-BNC, Syskiyou) was sharpened in 0.5 N NaOH with a microelectrode etcher (EE-1D, Bak Electronics) at 24V for 9 cycles. No more than three sensilla from the same antenna were recorded.

AC signals (100-20k Hz) and DC signals were simultaneously recorded on an NPI EXT-02F amplifier (ALA Scientific Instruments) and digitized at 5 kHz with Digidata 1550 (Molecular Devices). ORN spikes were detected and sorted using threshold search under Event Detection in Clampfit 10.4 (Molecular Devices). Spike timing data were exported and analyzed in Igor Pro 6.3 (Wavemetrics). Peri-stimulus time histograms (PSTHs) were obtained by averaging spike activities in 50-ms bins and smoothed using a binomial filter (Igor Pro 6.3, Wavemetrics).

Sensillum types were identified based on their locations on the antenna or maxillary palp, and their characteristic odor response profiles^14,20^. For ac3I and ac3II sensilla, sensillum types were determined according to ORN-specific fluorescent labeling (ac3I: *Ir75b-GAL4*; ac3II: *Ir75c-GAL4*) because the response profiles for ac3I and ac3II in *D. melanogaster* are virtually indistinguishable^26^. Based on the location of the fluorescence signals, we recorded from ac3I with a medial mounting position and ac3II with a posterior mounting position^52^.

**Odor stimuli**. Chemicals were >99% pure or of the highest purity available at Sigma-Aldrich unless otherwise specified. Odorants were diluted in paraffin oil unless otherwise noted. Apple cider vinegar (Spectrum, naturals filtered apple cider vinegar) and geosmin were diluted in water, and *trans*-palmitoleic acid (Cayman Chemical) was diluted in ethanol. For odor mixture experiments, individual odorants (2x stock solutions) were mixed either with paraffin oil or with another odorant at 1:1 (v/v) ratio prior to experiments. For short odor pulses, odor stimuli (100 μl applied to a filter disc) were delivered from a Pasteur pipette via a pulse of air (200 ml/min) into the main air stream (2000 ml/min). A Pasteur pipette filled with pure CO_2_ was used to deliver CO_2_ stimuli, as described previously^10^. Background odor stimuli were delivered from a 125-ml flask containing 3 ml of odor dilutions directly downstream of the main air stream (2000 ml/min).

For palmitoleic acid, 4.5 μl of the freshly diluted odorant was applied to filter paper inserted inside a truncated 200-µl pipette tip. Ethanol was allowed to evaporate for 1 hour in a vacuum desiccator prior to experiments. The odor cartridge was positioned around 4 mm away from the antenna as described^51^. Odor stimulus was delivered via a 500-ms pulse of air (500 ml/min) directly at the antenna in the presence of humidified air flow at 2000 ml/min from a different direction. Of note, female at4A does not respond to palmitoleic acid as strongly as male at4A in 7-day old flies (Ng et al., unpublished data).

**Optogenetic stimulation**. Newly eclosed female flies expressing the H134R-ChR2 transgene in targeted ORNs were reared in constant darkness for 5 days on fly food supplemented with 100 μM all *trans*-retinal (Sigma) unless otherwise specified. Flies were transferred to fresh retinal food one day prior to experiments. A light stimulus was generated via a blue LED (470 nm, Universal LED Illumination System, pE-4000, Cool LED). Light pulses (500-ms duration) were controlled by a shutter (Vincent Associates) driven by Clampex 10.4 (Molecular Devices). Light output around the position of the recorded antenna was measured with an optical power meter (PMKIT-05-01, Newport Corporation) via a slim profile wand detector (818-ST2/DB, Newport Corporation).

The LFP light responses in the ac3 neurons expressing H134R-ChR2 were too small to be reliably analyzed. Attempts to express H134R-ChR2 in ab5A (Or82a-GAL4) or ab5B (Or47a-GAL4) failed to yield any light responses, despite the fact that the fluorescence of mCherry tag on H134-ChR2 was visible in the target ORNs. Aging the transgenic flies to 14 days old or increasing retinal concentrations in the food did not improve the situation.

**Immunohistochemistry**. 7-day old female files expressing myc-tagged Orco and/or Or85a odorant receptors in the ab2B ORNs were anesthetized on ice, with their heads aligned in a collar, covered with Cryo-OCT (Tissue-Tek, Fisher Scientific), and frozen on dry ice as described^53^. Cryosections (14-μm) were fixed with 4% paraformaldehyde in 1X phosphate-buffered saline (PBS) and stained with rabbit anti-myc antibody (1: 250, 71D10, Cell Signaling Technology) and 21A6 (ciliary base marker, 1:200, DSHB), followed by goat anti-rabbit Alexa 647 secondary antibody (1: 250, A21236, Life Technologies) and goat anti-mouse Alexa 568 (1:200, #11019, Life Technologies). Confocal microscopy was performed with a Zeiss 880 Airyscan Microscope, and images were processed with ImageJ software.

**Sample preparation for SBEM**. Target expression of APEX2 in ORNs for SBEM was performed as described^44^. Briefly, transgenic *Drosophila* lines (*10xUAS-myc-APEX2-Orco* or *10xUAS-mCD8GF-APEX2*) were generated to facilitate dendritic targeting of APEX2^44^. Expression of APEX2 in select ORNs was driven by OrX-GAL4 drivers (Supplementary Table 3). Six to eight days old female flies were cold anesthetized prior to the dissection of their antennae. The antennae were processed with the CryoChem method^44^, which involves cryofixation by high-pressure freezing, freeze-substitution, rehydration, DAB labeling reaction, *en bloc* heavy metal staining, dehydration and resin infiltration.

Microcomputed X-ray tomography was performed on resin-embedded specimens using a Versa 510 X-ray microscope (Zeiss) to determine DAB-labeled region of interest. The specimens were then mounted on aluminum pins with conductive silver epoxy (Ted Pella) and sputter coated with gold-palladium for SBEM imaging. The ab3, ab4, ac3II, at4 datasets were collected with a Gemini SEM (Zeiss) equipped with a 3View block-face unit (Gatan); the ab5 dataset was collected with a Merlin scanning electron microscope (Zeiss) equipped with a 3View2XP and OnPoint backscatter detector (Gatan). Parameters for SBEM image acquisition are listed in Supplementary Table 5.

**Segmentation of DAB-labeled *Drosophila* ORNs**. The DAB-labeled *Drosophila* ORN was segmented in a semi-automated fashion using the IMOD software^54^ to generate a 3D model as described^44^. The IMOD command line ‘imodauto’ was used for the auto-segmentation by setting thresholds to isolate the labeled neuron of interest. Auto-segmentation was followed by manual proofreading and correction of errors by two independent proofreaders. The neighboring, unlabeled ORN was manually segmented using the same software. Due to insufficient DAB-labeling, the fine outer dendritic branches of most basiconic ORNs could not be reliably identified for segmentation.

**Morphometric analysis**. The 3D model of each ORN was first separated into cell body, inner dendrite and outer (sensory) dendrite models. The inner and outer dendrites were separated at the cilium base, a notably constricted dendritic region^40^. The volume measurements of ORNs were then obtained with the “imodinfo” function in IMOD based on the 3D models.

The lengths of most inner and outer dendrites were determined by first converting the 3D models into binary image files using the IMOD command “imodmop”. Then the skeletons of the 3D images were extracted using Skeletonize3D (https://imagej.net/Skeletonize3D) plugin in Fiji (NIH). The lengths of the resulting skeletons were obtained by the Fiji “Analyze Skeleton” function. For the ab3A, ab4A and ab5A ORNs, which exhibit significant dendritic enlargements (Fig. 6f), their inner dendritic lengths were determined by first visually identifying the center point in every ninth contour of the 3D models (300~400 nm z-step), then manually measuring and summing the distances between those points. The inner dendritic lengths of the ab3B, ab4B and ab5B ORNs were determined in the same way. The outer dendritic lengths reported in Supplementary Table 2 are the distances between the cilium bases and the tips of the longest dendritic branch.

The outer dendrites of ORNs were assumed to be cylindrical and their surface areas were calculated based on the measured volumes and lengths accordingly. The surface areas of cell bodies were measured with the “imodinfo” function in IMOD.

**Statistics**. All data presented as mean ± s.e.m were analyzed using Igor Pro 6.3 or SigmaPlot 13.0. Coefficients and the standard deviations of the linear fits were generated in Igor Pro 6.3. Unpaired two-tailed t-test was performed in Fig. 5 for single variable comparison between two groups. Paired two-tailed *t*-test was performed in Fig. 6 for the morphological comparison between grouped ORNs. Data are presented as mean ± s.e.m. *P* < 0.05 was considered to be statistically significant and is presented as ^∗^ *P* < 0.05, ^∗∗^ *P* < 0.01, or ^∗∗∗^ *P* < 0.001. For Fig. 4 and Supplemental Fig. 7, statistical significance for linear coefficients was determined by analysis of covariance (ANCOVA) in RStudio using functions within the car package (version3.0-0). Data are presented as coefficient ± s.d.. *P* < 0.05 was considered to be statistically significant and the differences are denoted by different letters.

## MODELING

### 1. Passive electric circuit model

We consider the passive electric circuit model containing an auxiliary cell (denoted “A”) and two ORNs (ORN_1_ & ORN_2_) (Fig. 7a). Each cell is modeled as an effective Thevenin circuit with battery and resistances. The auxiliary cell is modeled as a battery *E_A_* in line with an input resistance *R_A_*. The soma of ORN_1_ is modeled as a battery *E*_1_ with input resistance *R_in_*_1_, and the sensory dendrite is modeled as an odorant stimulation-dependent resistance *R_d_*_1_ (similar for ORN_2_). The driving forces for the currents are the batteries *E_A_*, *E*_1_ and *E*_2_. The voltage *V_A_* is the transepithelium potential, and *V_m_*_1_ and *V_m_*_2_ are the neuronal transmembrane potentials that control spike firing. Changes in *V_A_* are recorded as changes in the local field potential, LFP. The system of equations for the currents *I_A_*, *I*_1_ and *I*_2_ reads

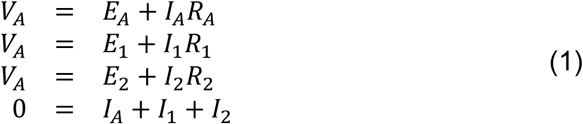

where we introduce the total resistances *R*_1_ = *R_in_*_1_ + *R_d_*_1_ and *R*_2_ = *R_in_*_2_ + *R_d_*_2_. By solving these equations, we obtain the currents

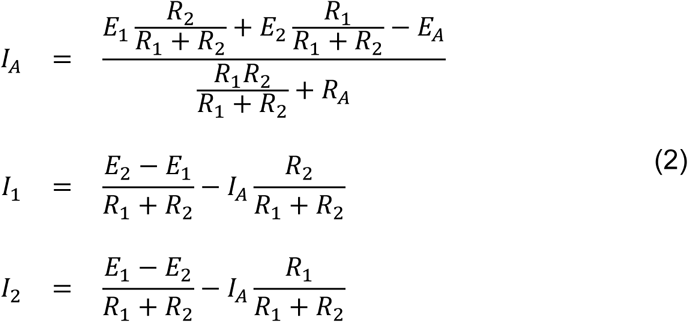

The neuronal part is an effective Thevenin circuit with battery 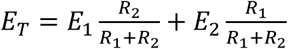 and sresistance 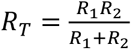. The neuronal membrane potentials are

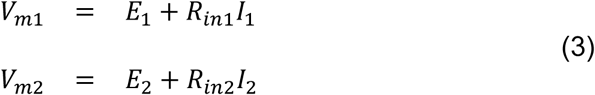

To further simplify the analysis, we use *R_A_* to define dimensionless resistances 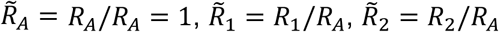 and rescale currents 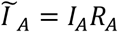, 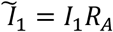and 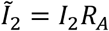. This does not affect the voltages in Eq. 1 and Eq. 3 and is formally equivalent to setting *R_A_* =1. In the following, we therefore omit the tilde symbols and simply use *R_A_* =1. As mentioned before, this does not affect the results and conclusions concerning voltages.

#### 1.1 Input and dendritic resistances

We are interested in how odorant stimulation and the morphometric properties of soma and sensory dendrites affect the transmembrane potentials *V_m_*_1_ and *V_m_*_2_. We assume that the input resistances *R_in_*_1_ and *R_in_*_2_ depend on the size of the soma and a smaller ORN has a higher input resistance compared to a larger ORN. Specifically, we assume that *R_in_*_1_ and *R_in_*_2_ are inversely proportional to the soma surfaces *A_S_*_1_ and *A_S_*_2_,

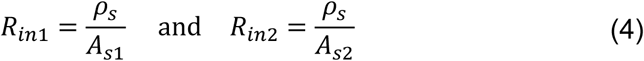

where *ρ_s_* is the soma membrane resistivity.

We model the sensory dendrite of an ORN as a uniform cylinder of length *L* with axial (cytoplasmic) resistivity *r_a_* and membrane resistivity *r_m_*, where *r_m_* depends on odor activation. From linear cable theory, the input resistance *R_s_* of a uniform leaky cylinder with an open and a sealed end is

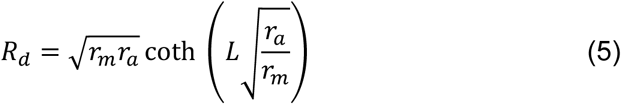

where 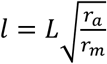 is the electrotonic length of the cylinder and 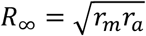 is the input resistance of an infinitely long cylinder. We further simplify Eq. 5 assuming that the cytoplasmic resistance is much smaller than the membrane resistance such that *l* ≪ 1 ^55^. By expanding Eq. 5 for *l* ≪ 1 we get in first order

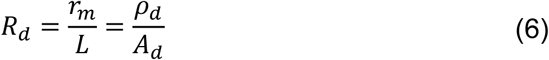

where *ρ_d_* is the dendritic membrane resistivity and *A_d_* is the dendritic surface.

The membrane resistivity *ρ_d_* implicitly depends on odor activation because odorants activate receptors that increase the membrane conductivity. We simplify the transduction process and assume that for a given stimulation with odorant concentration *od*, the amount of activated receptors is given by the Hill function

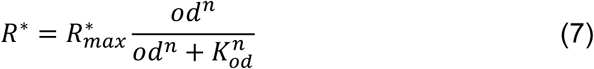

where *n* is the Hill-coefficient and *k_od_* the odorant concentration that activates half of the receptors. Receptor activation induces the additional membrane conductivity 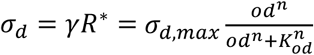. With the basal resistivity *ρ_d_*_,0_, the overall conductivity is 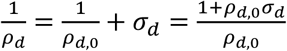. This gives

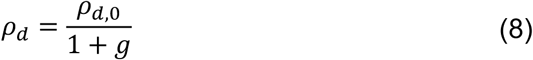

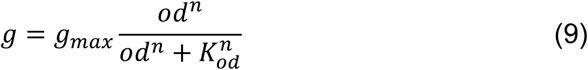

and 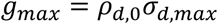.

In summary, to model the dendritic resistances we use the formula

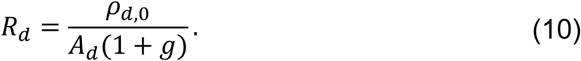

#### 1.2 Change in local field potential due to stimulation of ORN_1_ or ORN_2_

When the activation of ORN_1_ is altered due to odorant stimulation, the dendritic resistance *R_d_*_1_ changes by Δ*R_d_*_1_ which alters the local field potential LFP by |Δ*V_A_*|. From Eqs. 1~3 we compute that the corresponding changes in the membrane potentials are 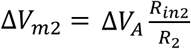 and

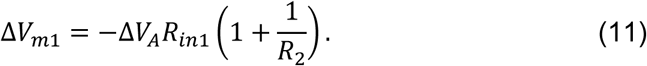

Similarly, when ORN_2_ is stimulated we have 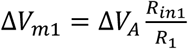 and

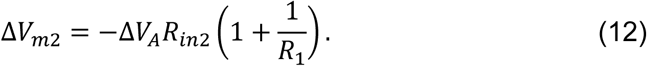

Because *R*_2_ (*R*_1_) remains constant when ORN_1_ (ORN_2_) is stimulated^11^, it follows that Δ*V_m_*_1_ and Δ*V_m_*_2_ change linearly with Δ*V_A_*. Assume that ORN_1_ corresponds to the larger neuron A and ORN_2_ to the smaller neuron B such that *R_in_*_1_ < *R_in_*_2_ and *R*_1_ < *R*_2_. With this we have 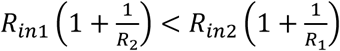 which is in agreement with Figure 3 showing that the slope of the spiking rate (which reflects Δ*V_m_*_1_ or Δ*V_m_*_2_) vs LFP is smaller for ORN_A_ compared to ORN_B_.

### 2. Fitting procedure

We are interested in whether the differences between our electrophysiological measurements can be explained by the morphometric differences among the grouped ORNs. To address this question, we focus on ab3, ab4 and ab5 sensilla for which we have electrophysiological and morphometric data. We simultaneously fitted the combined LFP data (Fig. 2) using a minimal model with a common set of basic parameters and ORN-specific parameters (surface area and odorant sensitivity). In the following, we derive the minimal model that we use for the fitting procedure. For values of fitting parameters, see section 2.4. The fitting and simulation results are presented in Fig. 7 of the main text.

#### 2.1 Estimating the batteries *E*_1_ and *E*_2_ using the resting membrane potential

We assume that the auxiliary cell is the same for all sensillum types, and therefore use a single value for *E_A_*. For *E*_1_ and *E*_2_, we do not assume that they are identical for all sensilla because ORN basal resistances differ depending on the morphometric feature of the neuron, and using the same *E*_1_ and *E*_2_ for all sensilla would lead to different resting membrane potentials for the ORNs (data not shown). This is problematic if one assumes that the mechanism of action potential generation is similar in all ORNs. Instead, we assume that at rest without odorant stimulation (basal condition) the membrane potentials of the ORNs are identical such that *V_m_*_1_ = *V_m_*_2_ = *V*_0_ with *V*_0_ = −60*_m_V*. We used the conditions *V_m_*_1_ = *V_m_*_2_ = *V*_0_ to express the parameters *E*_1_ and *E*_2_ as a function of *V*_0_. With Eqs. 2 and 3 we find

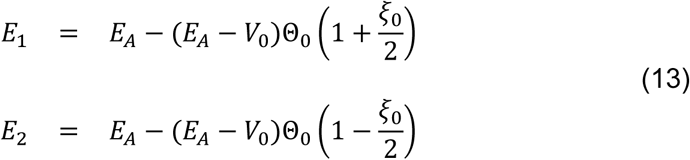

and

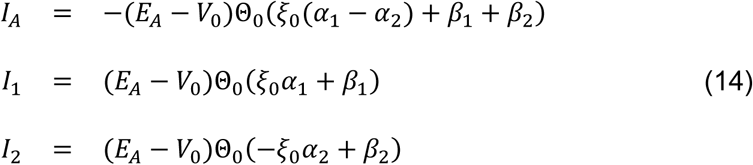

With

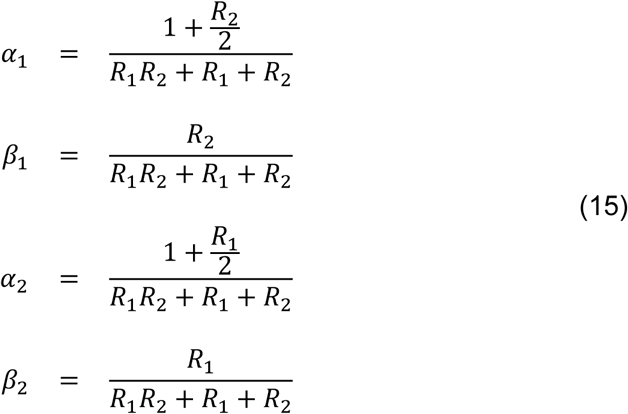

and

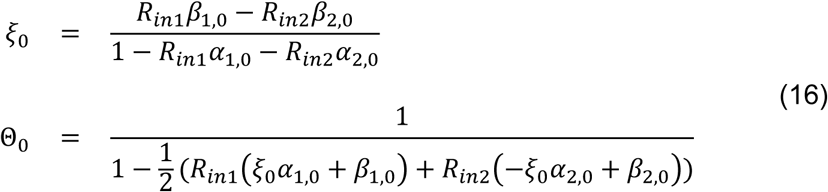

The values *β*_1,0_, *β*_2,0_, *α*_1,0_ and *α*_2,0_ are obtained from Eq. 15 by inserting basal dendritic resistances that depend on the neuronal morphometry. Eq. 14 shows that the effective driving force for the currents is *E_A_*−*V*_0_.

#### 2.2 Constraining the parameter space using the spike/LFP ratio

For the ab5 sensilla, we use the measured values for the outer dendritic surfaces *_Ad_*_1_ and *_Ad_*_2_. In contrast, for ab3 and ab4, the surfaces *_Ad_*_1_ and *_Ad_*_2_ are unavailable and are therefore free fitting parameters. However, for these sensilla, we use the measured spike/LFP ratio *r*(Fig. 3) to further constrain the parameter space. Our passive electrical model does not predict spiking rates. However, as discussed in section 1.2, we assume that Δ*V_m_*_1_ /Δ*V_A_* or Δ*V_m_*_2_/Δ*V_A_* is correlated to the spike/LFP ratio shown in Figure 3. Let *r*>1 be the ratio of the spike/LFP slopes of ORNA to ORNB shown in Figure 3. With the convention that ORN_1_ corresponds to ORNA and ORN_2_ to ORNB, we have

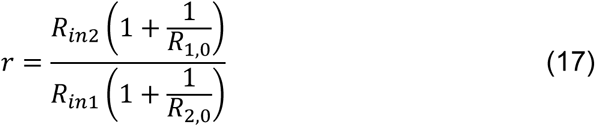

where *R*_1,0_ and *R*_2,0_ are basal resistances without odorant stimulation. We can use Eq. 17 to express *R*_1,0_ as a function of *R*_2,0_

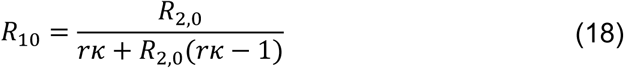

where 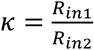. With 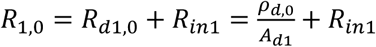 the dendritic surface of ORN_1_ is

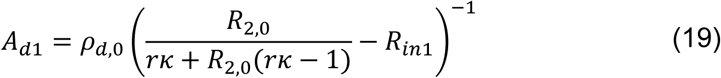

In summary, for ab3 and ab4, we use *_Ad_*_2_ and *r*as fitting parameters and *_Ad_*_1_ is computed with Eq. 19. The range for *r*is constrained by the values shown in Figure 3. In contrast, for ab5 we use our measured values for *_Ad_*_1_ and *_Ad_*_2_.

#### 2.3 Odorant dilution

For the odorant concentration, we write *od* = *c*_0_10*_m_*, where *x* < 0 corresponds to the odorant dilution and *c*_0_ is the initial undiluted concentration. By writing 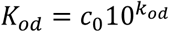 we have

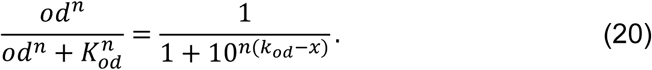

The parameter *K_od_* (resp. *K_od_*) characterizes the sensitivity of the ORN to the odorant and is specific for each ORN. We therefore allow for each neuron a different parameter *K_od_*.

#### 2.4 Parameters

(1) ORN-specific morphometric parameters. Measured input parameters are in black (see also Supplementary Table 2), and fitted values in blue.

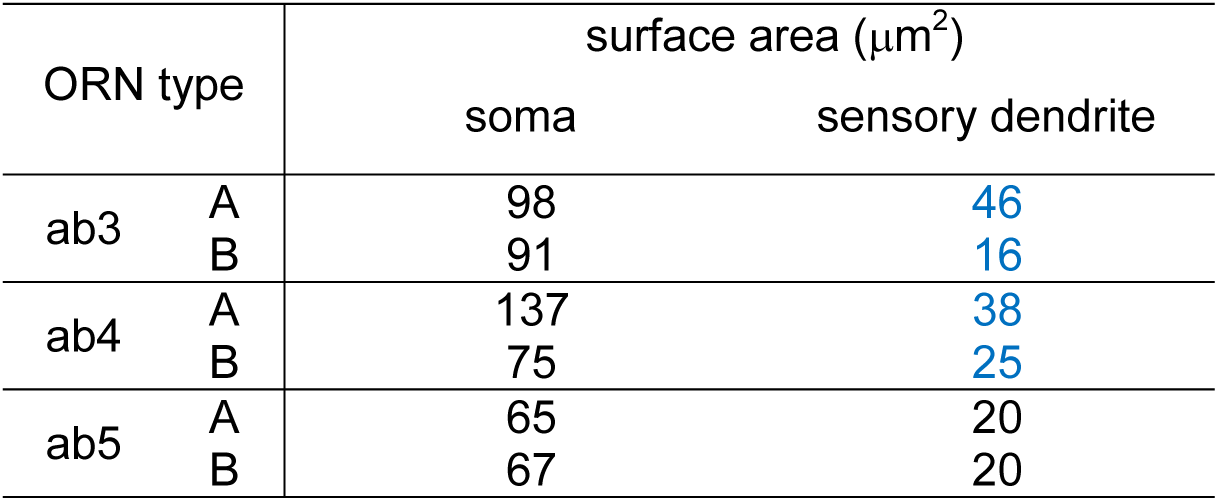

| ORN type | | surface area ($\mu\text{m}^2$) | |
| --- | --- | --- | --- |
|  |  | soma | sensory dendrite |
| ab3 | A | 98 | 46 |
|  | B | 91 | 16 |
| ab4 | A | 137 | 38 |
|  | B | 75 | 25 |
| ab5 | A | 65 | 20 |
|  | B | 67 | 20 |

(2) ORN-specific odorant sensitivities

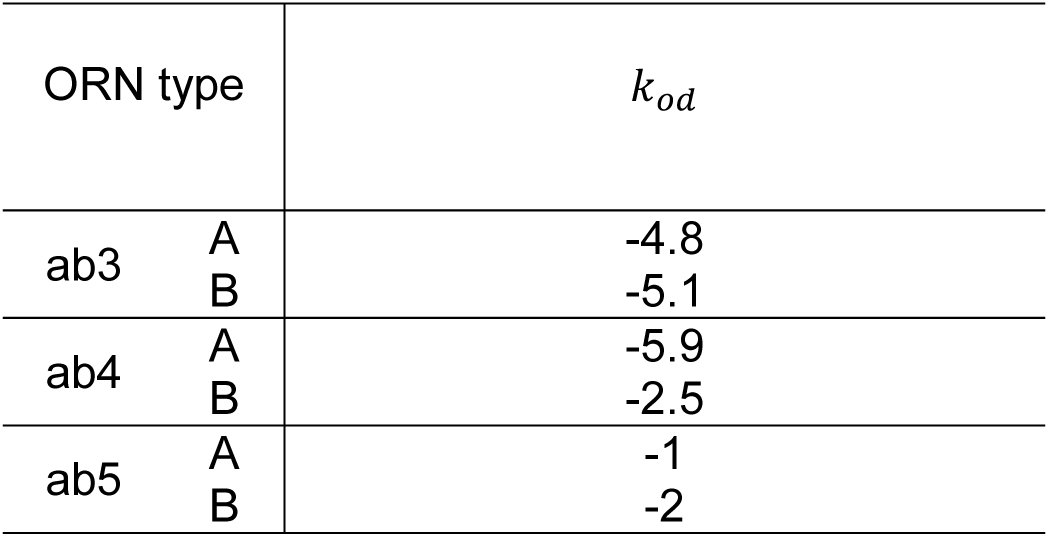

(3) Common parameters

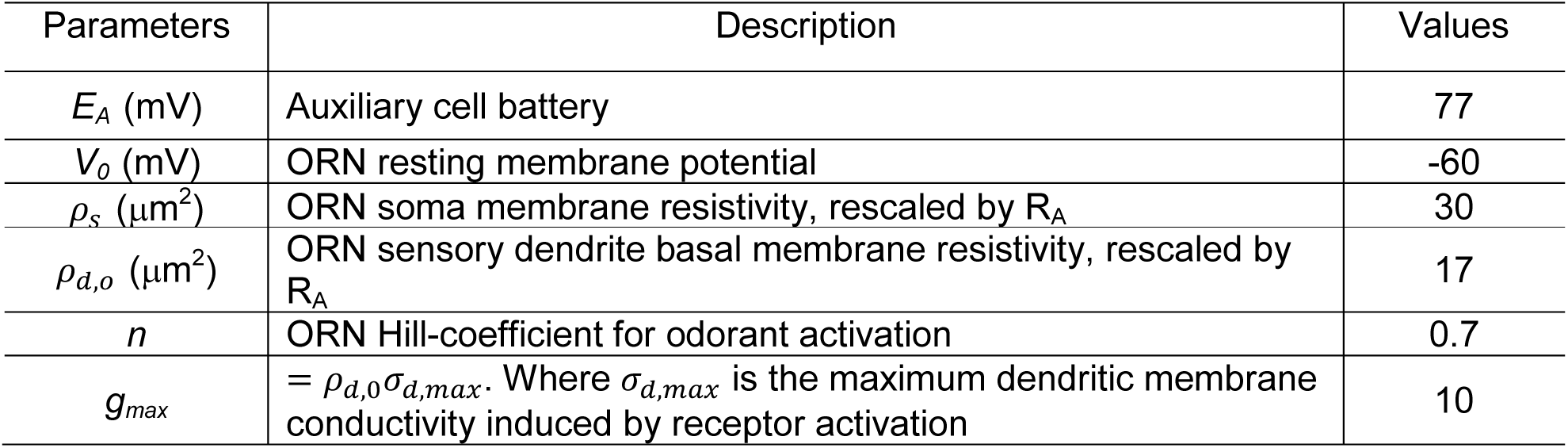

**Supplementary Figure 1.**
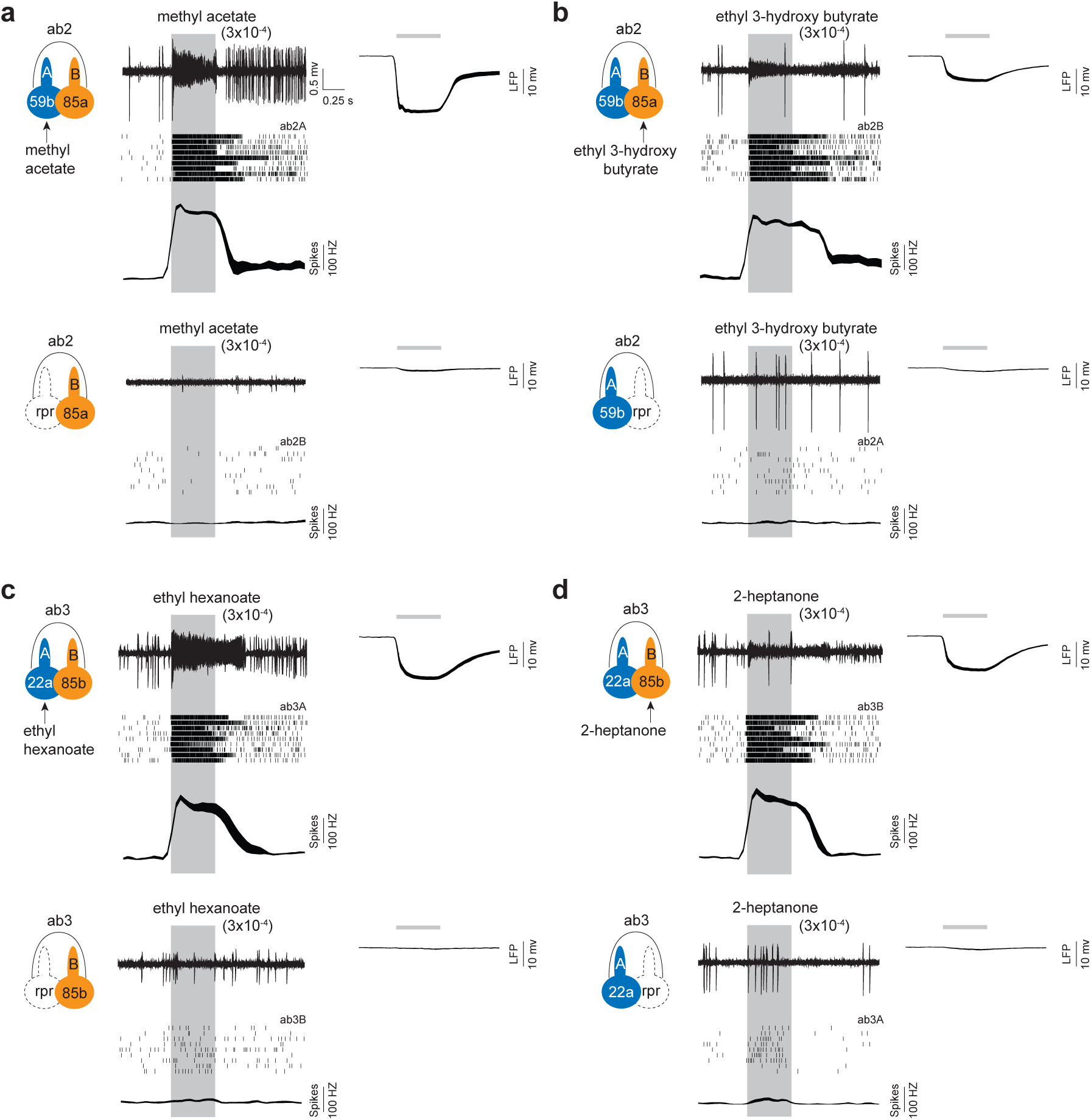
Identification of private odorants for grouped ORNs. Odorants from published datasets were selected to screen for private odorants^20,26,39,56–59^. **(a)** Top panels: ab2A is strongly activated by methyl acetate (3x10^-4^ dilution). Sample trace for the spike response (top), ab2A raster plots (middle), and the average ab2A peri-stimulus time histogram (bottom) are shown. Upper right panel: the corresponding LFP response. Line width indicates s.e.m. Bottom panels: a cell death gene, reaper (rpr) is expressed in ab2A to selectively ablate the neuron. In the absence of ab2A, methyl acetate (3×10^-4^) scarcely elicits any LFP response in the ab2 sensillum, indicating that methyl acetate-elicited LFP responses originate mainly from ab2A activation. *n=9*, mean ± s.e.m., parallel experiments. (**b**) As in **(a)**, except that ab2B responses are shown. Ethyl 3-hydroxy butyrate is a private odorant for ab2B. Genetic ablation of ab2B abolishes the LFP response elicited by ethyl 3-hydroxy butyrate (3x10^-4^ dilution) in the ab2 sensillum. **(c-d)** Identification of private odorants for ab3A **(c)** and ab3B **(d)**. Single-sensillum recordings and genetic ablation experiments were carried out as described in **(a)**. Private odorants do not elicit significant LFP responses in the absence of the target ORNs. The odorants are not considered “private” above the indicated concentrations because they will also activate the neighboring neurons. *n=9*, mean ± s.e.m., parallel experiments. Similar genetic ablation experiments were conducted to identify private odorants for 7 additional pairs of grouped ORNs (see **Supplementary Table 1** for summary).

**Supplementary Figure 2.**
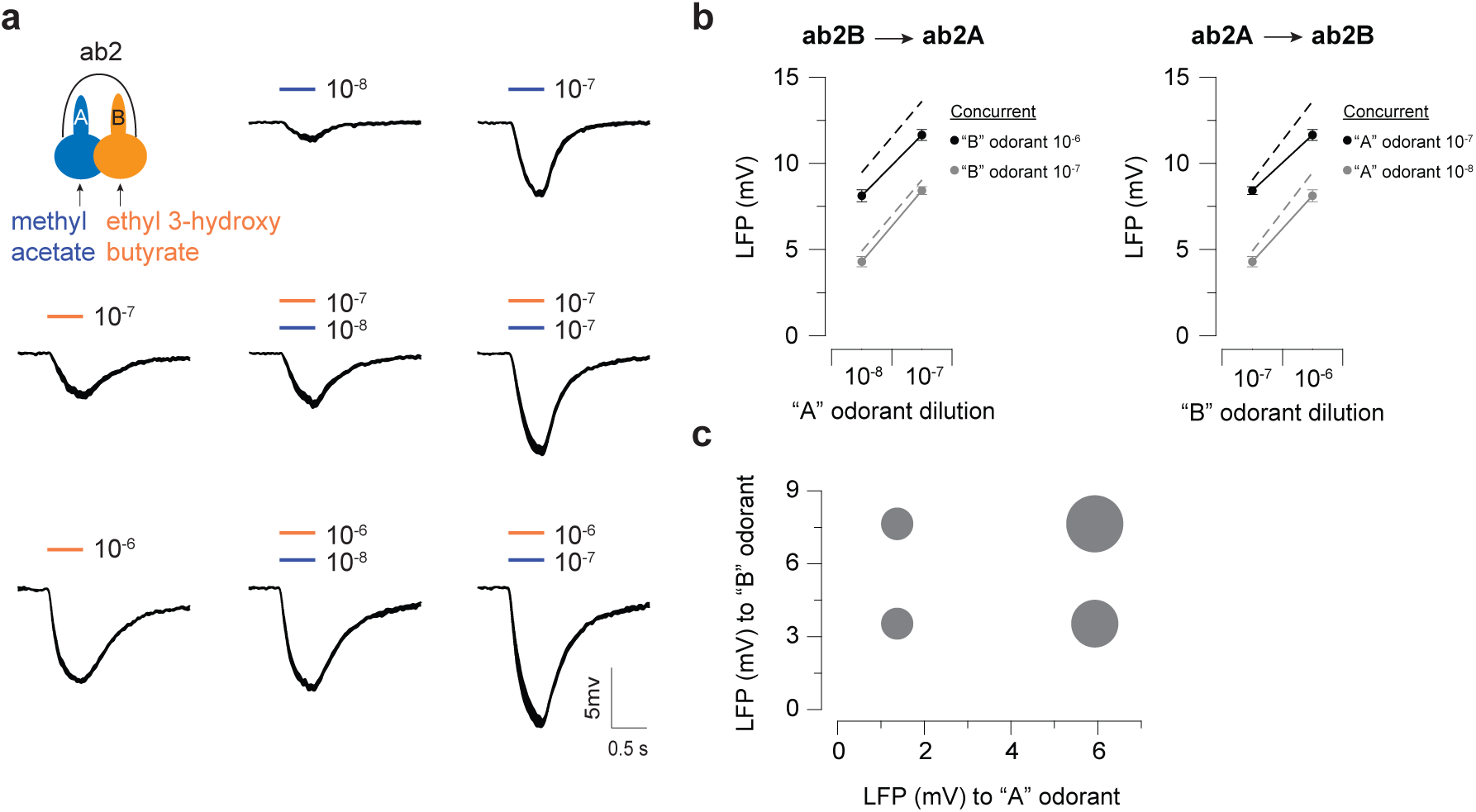
The degree of ephaptic inhibition is influenced by the field responses of grouped ORNs. **(a)** LFP responses of the ab2 ORNs to 0.5-sec pulses of their private odorants (ab2A: methyl acetate, blue bar; ab2B: ethyl 3-hydroxy butyrate, orange bar). Odorants were delivered either as individuals or as concurrent binary odor mixtures. *n=8* sensilla, line width indicates s.e.m. **(b)** Peak LFP responses (absolute values) are plotted as a function of odorant dilution for methyl acetate (left panel) or ethyl 3-hydroxy butyrate (right panel). The concentrations of the concurrent odorants that activate the neighboring neurons are indicated. The linear sums of the LFP responses to individual private odorants are connected by dashed lines, predicting the LFP responses to odor mixtures if there is no ephaptic inhibition. The measured LFP responses were smaller than the linear sums, indicating ephaptic inhibition between ORNs, mean ± s.e.m. **(c)** Bubble plot of the magnitude of inhibition in relation to the LFP responses to methyl acetate (“A” odorant, x-axis) or ethyl 3-hydroxy butyrate (“B” odorant, y-axis). Inhibition was determined by subtracting the measured LFP response from the linear sum. The size of the bubble scales with the magnitude of inhibition.

**Supplementary Figure 3.**
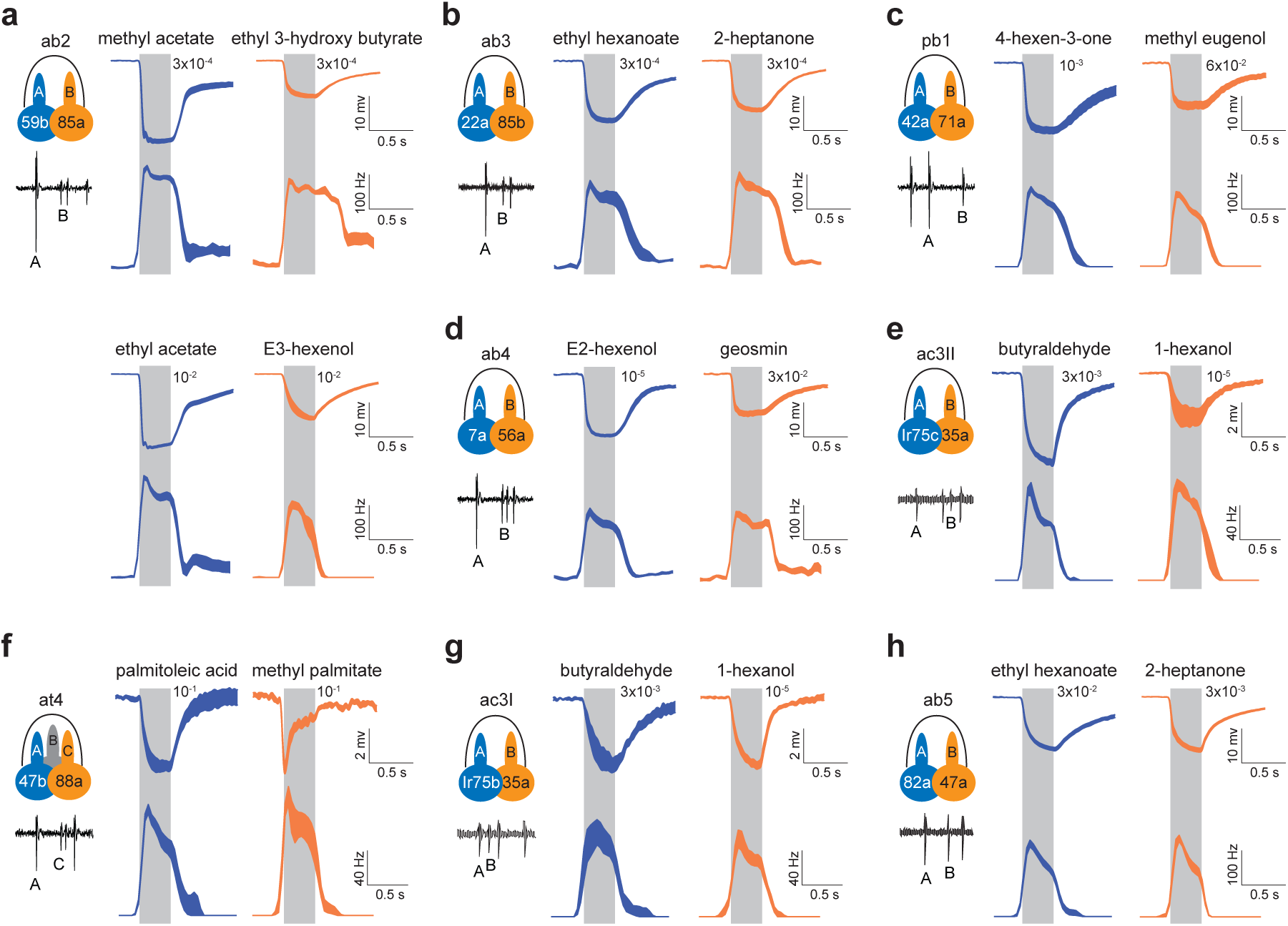
LFP and the corresponding spike responses of grouped ORNs to private odorants. ORNs were stimulated with their respective private odorants at the highest possible concentrations (see also **Supplementary Table 1**). Average LFP and the corresponding spike responses are shown for the large-spiking “A” neurons (blue traces) and the small-spiking neighbors (orange traces). Gray rectangles denote periods of odor stimulation (0.5 sec). The concentrations of each odorant are indicated. **(a-h)** Responses we¬¬re record-ed from neighboring neurons housed in the same sensillum. Eight sensillum types were examined (ab2, ab3, ab4, pb1, ac3 type I, ac3 type II, at4). *n=9* pairs of ORNs, except for ac3: *n=6* pairs, line width indicates s.e.m.

**Supplementary Figure 4.**
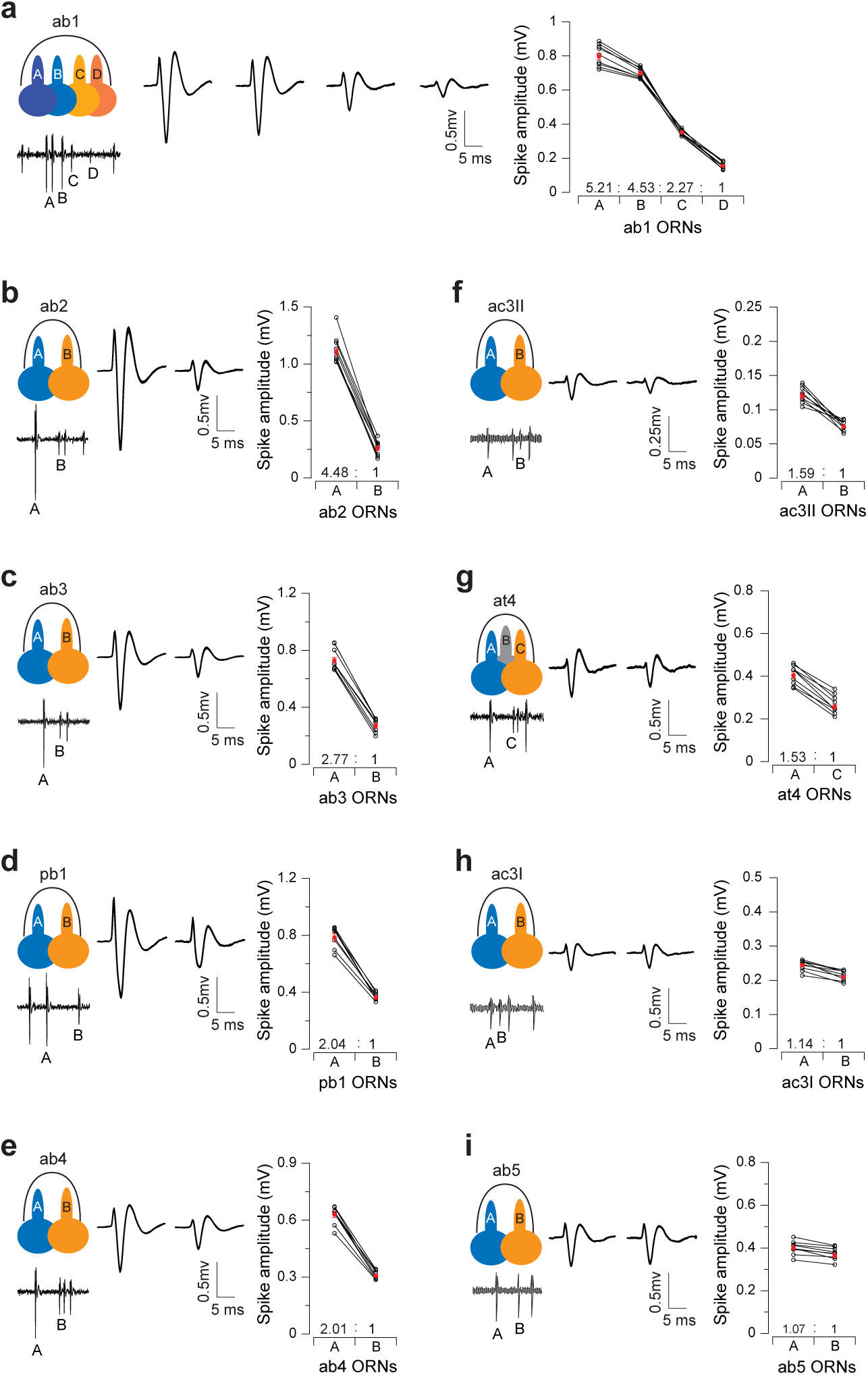
Relative extracellular spike amplitudes of grouped ORNs. Spontaneous spike activities from compartmentalized ORNs were recorded to characterize their extracellular spike amplitudes in nine different types of sensilla **(a-i)**. Recordings were bandpass filtered at 100-20k Hz. Left: Average spike waveforms are shown for each of the grouped ORNs. Line width indicates s.e.m. Right: Comparison of the spike amplitudes of compartmentalized ORNs. Each data point represents the average spike amplitude of an ORN based on its spontaneous activity. Lines connect measurements from the same recording. Red dots denote average spike amplitudes. Spike amplitude ratios, relative to the paired ORN with the smallest spike, are indicated for each sensillum types. *n=9*, mean ± s.e.m. Note: for technical issues, the signal-to-noise ratio for ac3 type II is lower than that for other sensillum types and the absolute spike amplitudes may thus be underestimated. See Online **Methods** for the identification of ac3 type I and type II sensilla.

**Supplementary Figure 5.**
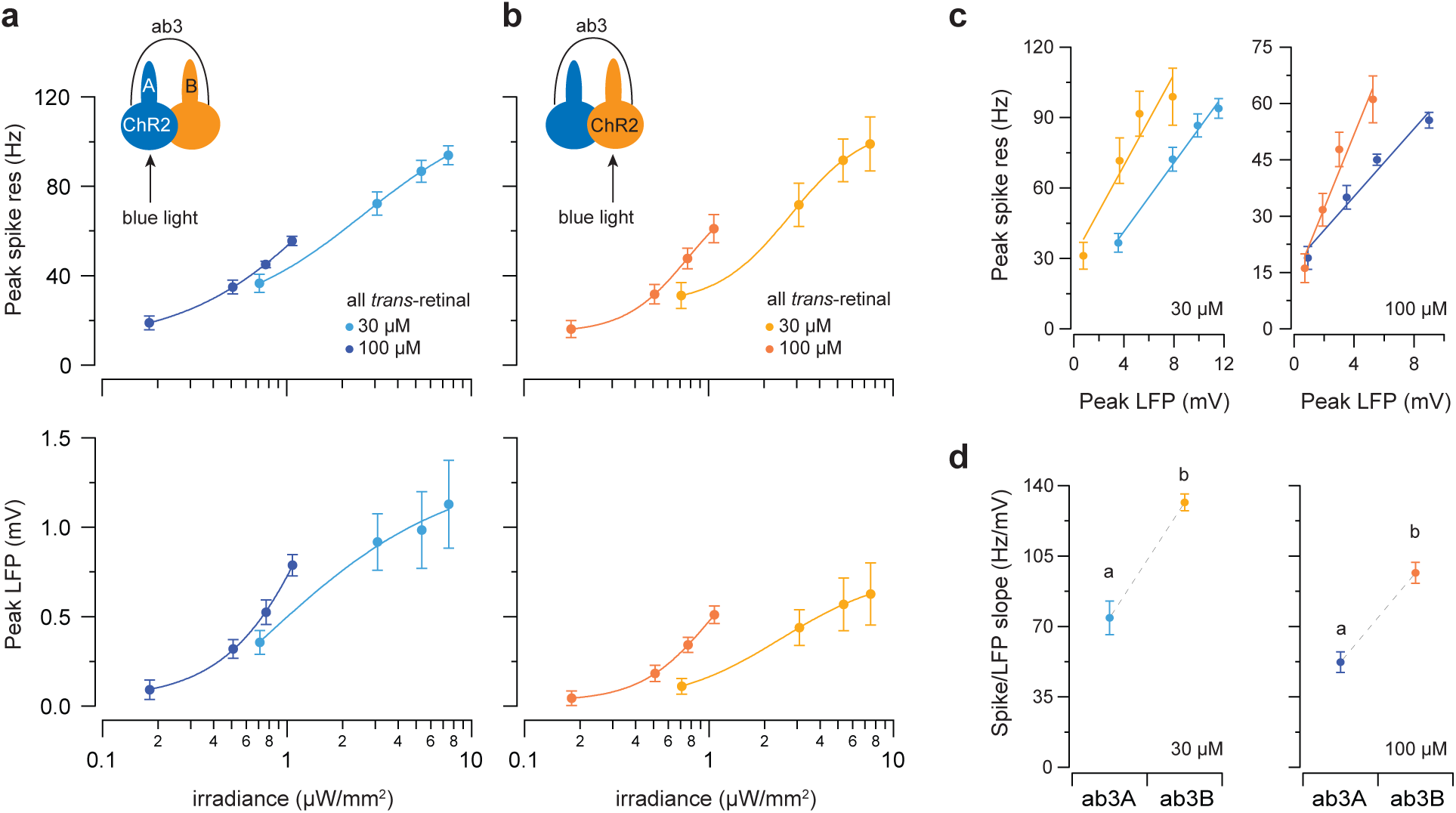
Optogenetic analysis with different retinal concentrations. **(a-b)** H134R-Channelrhodopsin2 (ChR2) was expressed in either ab3A **(a)** or ab3B **(b)** by the GAL4-UAS system. Newly emerged female flies were fed with the chromophore, all trans-retinal, at indicated concentrations (30 or 100 μM) for 5 days prior to experiments. ORNs were activated by 500-ms pulses of blue light of graded irradiances. Dose-response curves are shown to demonstrate the peak spike (top panels) and peak LFP responses (bottom panels). Results are from parallel experiments for each retinal concentration. *n=9*, mean ± s.e.m. (**c**) Peak spike responses are plotted as a function of peak LFP responses. Lines indicate linear fits *(y = ax + b)*. The spike/LFP relationships are shown based on the retinal concentrations. (**d**) The respective “a” coefficients (spike/LFP slope) for ab3A and ab3B are plotted for comparison. Dotted lines link results from parallel experiments. Statistical analysis was performed with ANCOVA and significant differences (*P* < 0.05) are denoted by different letters. Error bars = s.d.

**Supplementary Figure 6.**
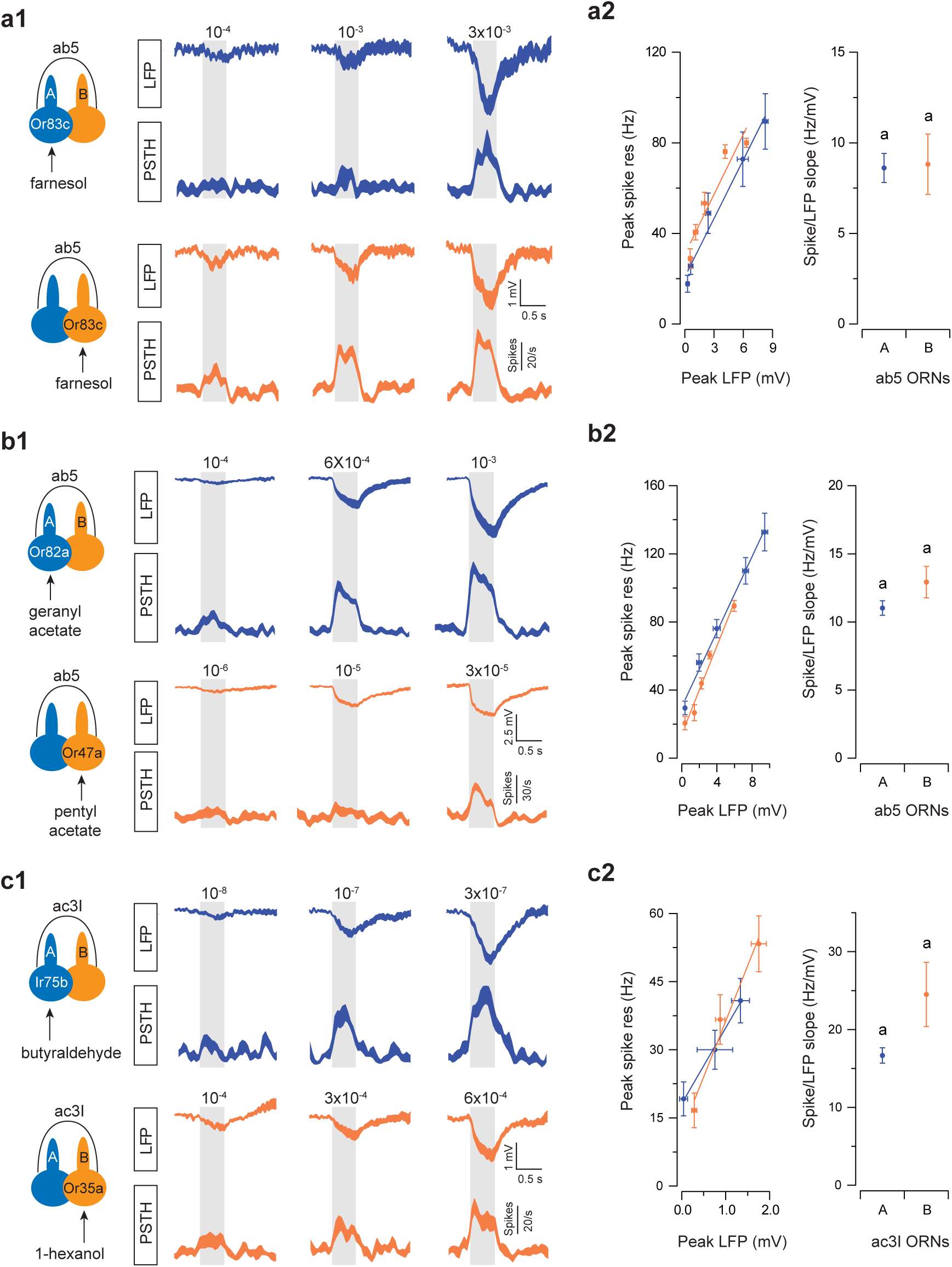
Spike-LFP analysis in grouped ORNs of similar spike amplitudes. (a) Or83c was ectopically expressed in either ab5A or ab5B using the GAL4-UAS system. ORNs were selectively activated by the odorant, farnesol, at different concentrations. **(a1)** Average LFP responses and the corresponding spike responses are shown for ab5A (blue traces) and ab5B ORNs (orange traces). **(a2)** Peak spike responses are plotted as a function of peak LFP responses. Left panel: Lines indicate linear fits *(y = ax + b)*. *n=9*, mean ± s.e.m. Right panel: The respective “*a*” coefficients (spike/LFP slope) for ab5A and ab5B are plotted for comparison. Error bars = s.d. **(b-c)** Similar to **(a)** except that ab5 ORNs **(b)** or ac3I ORNs **(c)** were selectively activated by their cognate private odorants at different concentrations. *n=9* pairs of ORNs for **(b)** and 6 pairs for **(c)**, mean ± s.e.m. Statistical analysis was performed with ANCOVA and significant differences (*P* < 0.05) are denoted by different letters.

**Supplementary Figure 7.**
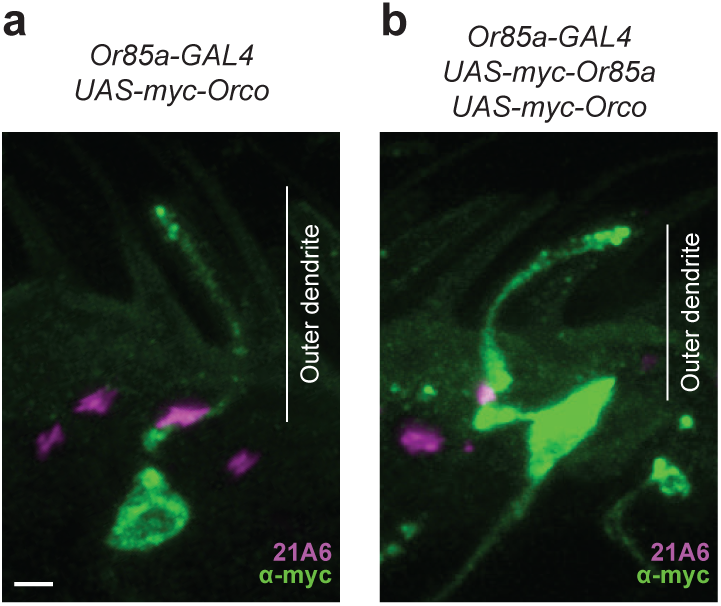
Dendritic targeting of ectopically expressed odorant receptors in ORNs. Confocal images of antennal sections immunolabeled with an anti-myc antibody (green) and an antibody specific for the sensory cilium base marker 21A6 (magenta). Ectopic expression of myc-tagged Orco **(a)** or myc-Orco and myc-Or85a **(b)** in the ab2B ORNs using Or85a-GAL4 driver. Immunofluorescence signal of myc labeling is observed throughout the target ORNs, including the sensory dendrites where olfactory transduction takes place. The anti-myc signal was stronger when myc-ORCO and myc-OR85a were co-expressed. Images in **(a)** and **(b)** were acquired in parallel with identical parameters. Scale bar, 2 μm.

